# Renal Proximal Tubule Cell-specific Megalin Deletion Does Not Affect Atherosclerosis But Induces Tubulointerstitial Nephritis in Mice Fed Western Diet

**DOI:** 10.1101/2024.05.11.592234

**Authors:** Naofumi Amioka, Michael K. Franklin, Masayoshi Kukida, Liyuan Zhu, Jessica J. Moorleghen, Deborah A. Howatt, Yuriko Katsumata, Adam E. Mullick, Motoko Yanagita, Michelle M. Martinez-Irizarry, Ruben M. Sandoval, Kenneth W. Dunn, Hisashi Sawada, Alan Daugherty, Hong S. Lu

**Author notes:** These authors contributed equally (Naofumi Amioka, Michael K. Franklin, Masayoshi Kukida). **Corresponding Authors:** Hisashi Sawada Alan Daugherty Hong S. Lu.

## Abstract

**Background:** Pharmacological inhibition of megalin (also known as low-density lipoprotein receptor-related protein 2: LRP2) attenuates atherosclerosis in hypercholesterolemic mice. Since megalin is abundant in renal proximal tubule cells (PTCs), the purpose of this study was to determine whether PTC-specific deletion of megalin reduces hypercholesterolemia-induced atherosclerosis in mice.

**Methods:** Female *Lrp2* f/f mice were bred with male *Ndrg1*-*Cre ERT2* +/0 mice to develop PTC-LRP2 +/+ and −/− littermates. To study atherosclerosis, all mice were bred to an LDL receptor −/− background and fed a Western diet to induce atherosclerosis.

**Results:** PTC-specific megalin deletion did not attenuate atherosclerosis in LDL receptor −/− mice in either sex. Serendipitously, we discovered that PTC-specific megalin deletion led to interstitial infiltration of CD68+ cells and tubular atrophy. The pathology was only evident in male PTC-LRP2 −/− mice fed the Western diet, but not in mice fed a normal laboratory diet. Renal pathologies were also observed in male PTC-LRP2 −/− mice in an LDL receptor +/+ background fed the same Western diet, demonstrating that the renal pathologies were dependent on diet and not hypercholesterolemia. In contrast, female PTC-LRP2 −/− mice had no apparent renal pathologies. In vivo multiphoton microscopy demonstrated that PTC-specific megalin deletion dramatically diminished albumin accumulation in PTCs within 10 days of Western diet feeding. RNA sequencing analyses demonstrated the upregulation of inflammation-related pathways in kidney.

**Conclusions:** PTC-specific megalin deletion does not affect atherosclerosis, but leads to tubulointerstitial nephritis in mice fed Western diet, with severe pathologies in male mice.

## INTRODUCTION

Megalin, also known as low-density lipoprotein receptor-related protein 2 (LRP2), is a 520-kD transmembrane protein that belongs to the low-density lipoprotein (LDL) receptor family. During embryonic development, megalin plays a critical role in brain, cardiovascular, and lung development, as demonstrated by global megalin-deficient mice.^1–4^ Megalin becomes most abundant in renal proximal tubule cells (PTCs) after birth,^5^ and it functions primarily as an endocytic receptor in renal PTCs for many ligands including components of the renin-angiotensin system.

The renin-angiotensin system is important for blood pressure regulation and contributes to the pathogenesis of atherosclerosis.^6^ Angiotensin II (AngII) is the major effector in the renin-angiotensin system. Our previous study demonstrated that global inhibition of megalin by antisense oligonucleotides (ASO) administration attenuated hypercholesterolemia-induced atherosclerosis in both male and female mice, accompanied by diminished protein abundance of AGT and renin in renal PTCs as well as renal AngII concentrations.^7^ These findings support the hypothesis that megalin contributes to atherosclerosis by interacting with the renin-angiotensin components in PTCs.

To determine whether megalin in kidney contributes to atherosclerosis, PTC-LRP2 +/+ and −/− littermates were generated in an LDL receptor −/− background by breeding megalin floxed mice with transgenic mice expressing an inducible *Cre* driven by an N-myc downstream-regulated gene 1 (*Ndrg1*) promoter. In contrast to our findings in both male and female LDL receptor −/− mice administered *Lrp2* ASO,^7^ PTC-specific megalin deficiency did not attenuate hypercholesterolemia-induced atherosclerosis in either sex. Serendipitously, we found that megalin deficiency in PTCs led to tubulointerstitial (TIN) leukocyte infiltration and tubular atrophy predominantly in male PTC-LRP2 −/− mice with either LDL receptor +/+ or LDL receptor −/− background that were fed a Western diet.

## MATERIALS AND METHODS

### Data availability

Detailed materials and methods are available in this manuscript. Numerical data are available in the Supplemental Excel File. Bulk RNA sequencing data (raw FASTQ and aligned data) are publicly available at the gene expression omnibus repository (GEO accession number: GSE268879).

### Mice

*Lrp2* floxed (*Lrp2* f/f) mice were developed under a contract with the Ingenious Targeting Laboratory using the same strategy reported by Willnow and colleagues.^8^ Female *Lrp2* f/f mice were bred with male *Ndrg1*-*Cre ERT2* +/0 mice.^9^ The breeding strategy for generating *Ndrg1*-*Cre ERT2* 0/0 *Lrp2* f/f (PTC-LRP2 +/+) and *Ndrg1*-*Cre ERT2* +/0 *Lrp2* f/f littermates (PTC-LRP2 −/−) is shown in **Figure S1** and **Major Resources Tables**. To study atherosclerosis, these mice were bred further into LDL receptor −/− background. For mice injected with *Lrp2* ASO, Gen 2.5 ASOs at 6 mg/kg body weight dissolved in sterile PBS were administered via subcutaneous injections once a week for 13 weeks. The injections started 1 week before Western diet feeding was initiated.

To study renal pathologies, mice were bred to either LDL receptor +/+ or LDL receptor −/− on a C57BL/6J background. DNA was extracted from tails or kidneys using Maxwell DNA purification kits (Cat # AS1120; Promega). *Cre* genotype was determined before weaning and confirmed post-termination. Deletion of *Lrp2* was confirmed in kidney using either PCR, qPCR, or immunostaining of megalin.

All mice were maintained in a barrier facility on a light:dark cycle of 14:10 hours and fed a normal laboratory diet after weaning. To promote *Cre* translocation, mice at 4-6 weeks of age were injected intraperitoneally with tamoxifen (150 mg/kg/day) for 5 consecutive days. Two weeks after the last injection of tamoxifen, mice were fed a diet containing saturated fat (milk fat 21% wt/wt) and cholesterol (0.2% wt/wt; Diet # TD.88137, Inotiv, termed “Western diet”) for 12 weeks to develop atherosclerosis or renal pathologies. In studies investigating the underlying mechanisms of tubulointerstitial nephritis (TIN), mice were fed this Western diet for 10 days, 2 weeks, or 12 weeks, depending on the experimental purpose.

Both male and female littermates were used for the experiments reported in this manuscript in accordance with the Statement from the ATVB Council.^10^ At termination, mice were euthanized using overdose of a ketamine and xylazine cocktail. All animal experiments in this study were performed according to protocols approved by the University of Kentucky (Protocol number 2018-2968) or Indiana University (Protocol number 21052 for intravital microscopy) Institutional Animal Care and Use Committee.

### Western blot analysis

Kidney samples were homogenized in T-PER buffer (Cat # 78510, Thermo Scientific) with a protease inhibitor cocktail (Cat # P8340, Sigma-Aldrich) using Kimble Kontes disposable Pellet Pestles (Cat # Z359971, DWK Life Science LLC.). Protein concentrations were determined using a BCA assay kit (Cat # 23225, Thermo Scientific). Equal masses of protein per sample (3 µg) were resolved by SDS-PAGE and transferred electrophoretically to PVDF membranes. After blocking, antibodies against the following proteins were used to probe membranes: Megalin (0.1 µg/ml, Cat # ab76969, abcam), cubilin (1 µg/ml, Cat # PA5-115063, Invitrogen), and GAPDH (0.1 µg/ml, Cat # 14C10, Cell Signaling Technology). Subsequently, membranes were incubated with goat anti-rabbit secondary antibody (0.3 µg/ml, Cat # PI-1000, Vector Laboratories). Immune complexes were visualized by chemiluminescence (Cat # 34080, Thermo Fisher Scientific) using a ChemiDoc (Cat # 12003154, Bio-Rad) and quantified using Image Lab software (v6.0.0, Bio-Rad).

### RNA isolation and quantitative PCR (qPCR)

Total RNA was extracted from kidneys using Maxwell® RSC simplyRNA Tissue Kits (Cat # AS1340; Promega) in accordance with the manufacturer’s protocol. Total RNA was transcribed reversely to cDNA using iScript™ cDNA Synthesis kits (Cat # 170-8891; Bio-Rad). Quantitative PCR was performed to quantify *Lrp2* mRNA abundance in the kidney using SsoFast™ EvaGreen® Supermix kits (Cat # 172-5204; Bio-Rad) on a Bio-Rad CFX96 cycler. Data were analyzed using ΔΔCt method and normalized by the geometric mean of 3 reference genes: *Actb*, *Gapdh*, and *Rplp2*.

### Immunostaining

Immunostaining was performed using Xmatrx® Infinity, an automated staining system (Cat #: AS4000RX; BioGenex), on paraffin-embedded sections. After fixation using paraformaldehyde (4% wt/vol), kidney samples were incubated in ethanol (70% vo/vol) for 24 hours, embedded into paraffin, and cut at a thickness of 4 µm. Subsequently, sections were deparaffinized using limonene (Cat # 183164; Millipore-Sigma) followed by 2 washes with absolute ethanol (Cat # HC-800-1GAL; Fisher Scientific), and 1 wash with automation buffer (Cat # GTX30931; GeneTex). Deparaffinized sections were incubated with H_2_O_2_ (1% vol/vol; Cat # H325-500; Fisher Scientific) for 10 minutes at room temperature and then antigen retrieval (Cat # HK547-XAK; BioGenex) for 20 minutes at 98 °C. Non-specific binding sites were blocked using goat serum (2.5 % vol/vol; Cat # MP-7451; Vector laboratories) for 20 minutes at room temperature. Sections were then incubated with rabbit anti-megalin antibody (Cat # ab76969; abcam) diluted in primary antibody diluent (Cat #: GTX28208; GeneTex) for 15 min at 40 °C, and rabbit anti-CD68 (E3O7V) antibody (Cat # 97778; Cell Signaling Technology), rat anti-CD45 antibody (Cat # 553076; BD Pharmingen), anti-CD3 antibody (Cat # ab16669; abcam), anti-CD19 antibody (Cat # 90176S; Cell Signaling Technology), or AGT (Cat # 28101; IBL-America) overnight at 4 °C. Goat anti-rabbit or Goat anti-rat IgG conjugated with horseradish peroxidase (30 minutes, Cat # MP-7451 & MP-7444; Vector Laboratories) was used as the secondary antibody. ImmPACT® NovaRed (Cat # SK4805; Vector) was used as a chromogen, and hematoxylin (Cat # 26043-05; Electron Microscopy Sciences) was used for counterstaining. Slides were coverslipped with mounting medium (Cat # H-5000; Vector). Three negative controls were used for immunostaining: (1) no primary and secondary antibodies, (2) secondary antibody only, and (3) nonimmune rabbit IgG to replace the primary antibody. Images were captured using Axioscan Z1 or 7.

### Hematoxylin and eosin staining

Paraffin-embedded kidney sections were stained with hematoxylin and eosin. After paraffin removal, sections were stained with Harris hematoxylin (Cat # 26041, EMS) for 5 minutes and then rinsed with continuous tap water for 5 minutes. Subsequently, sections were stained with eosin (Cat # ab246824, Abcam) for 4 minutes, rinsed with 100% ethanol, and allowed to air dry. Images were acquired using Zeiss Axioscans (Z1 or 7).

### Periodic acid-Schiff (PAS) staining and quantification of glomerular damages

PAS staining was performed using Periodic Acid Schiff Stain Kit (Cat # ab150680, abcam) according to the manufacturer’s protocol. Glomerular damages were quantified and meaned based on measurements in 20 glomeruli/kidney/mouse by three researchers using a scoring system based on the extent of lesions, including nodular sclerosis and mesangial expansion: 0 = no lesion; 1 = <25%; 2 = 25–50%; 3 = 50-75%; and 4 = >75% of a glomerulus.^11,12^ All quantifications were performed double-blindly regarding study groups.

### Picrosirius red (PSR) staining and quantification of renal fibrosis

Deparaffinized sections were incubated with picrosirius red and fast green solution: Direct Red 80 (0.1% vol/vol; 365548, Sigma-Aldrich), Fast Green FCF (0.1% wt/vol; F7258, Sigma-Aldrich) in saturated picric acid (1.3% wt/vol; P6744, Sigma-Aldrich), for 60 minutes at room temperature. The slides were then rinsed with acidified water (0.1 N hydrochloride acid, H2505, Aqua Solutions Inc), dehydrated through increasing concentration of ethanol, incubated with three changes of xylene, and coverslipped with DEPEX (50-247-470, Thermo Fisher Scientific). Collagen-positive areas within the regions of interest (ROIs) of picrosirius red sections were measured as a binary area fraction using the following color threshold: collagen: R: 185-246, G: 0-208, B: 0-213. All measurements were verified by a member of the laboratory who was blinded to the identification of study groups.

### Systolic blood pressure measurements

Systolic blood pressure was measured on conscious mice by a non-invasive tail-cuff system (BP-2000, Visitech) following our standard protocol.^13^ Data were collected at the same time each day for three consecutive days before the termination. Criteria for accepted data were systolic blood pressure between 70 and 200 mmHg and standard deviation < 30 mmHg for at least 5 successful recorded data/mouse/day. The mean systolic blood pressure of each mouse from the 3-day measurements was used for data analysis.

### Quantification of atherosclerosis

Atherosclerotic lesions were traced manually on the intimal surface area of the aorta with an *en face* method in accord with the AHA Statement ^14^ and our standard protocol.^15^ Briefly, the aorta was dissected and fixed in neutrally buffered formalin (10% vol/vol) overnight. The adventitial tissues were removed, and the intimal surface was exposed by longitudinal cuts. Subsequently, the aorta was pinned on a black wax surface, and *en face* images were captured by a digital camera (DS-Ri1; Nikon). Atherosclerotic lesions were traced manually on the images from the ascending aorta to the descending thoracic aorta that was 1 mm distal from the left subclavian artery using a Nikon NIS-Elements software (NIS-Elements AR 5.11.0.). Raw data were verified independently by a senior staff member who was blinded to the identity of the study groups. Atherosclerotic lesion size was presented as percent lesion area showing below:

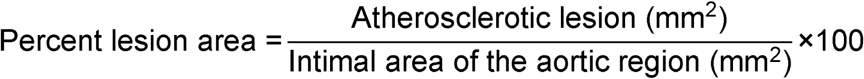

### Plasma total cholesterol concentrations

Mouse blood samples were collected with EDTA (1.8 mg/ml) via right ventricle at termination and centrifuged at 3000 g for 10 minutes at 4°C to collect plasma. Plasma total cholesterol concentrations at termination were measured using an enzymatic kit (Cat # 999-02601; FUJIFILM or Cat # C7510-120; Pointe Scientific).

### Urinary proteomics and ELISA

Urine was collected using metabolic cages (TSE Systems) or LabSand (Braintree Scientific, Inc.). Proteomics were performed using the method detailed in the Supplemental Materials. Urine AGT (Cat # ab245718, abcam), RBP4 (Cat # AG-45A-0012YEK-KI01; AdipoGen), KIM-1 (Cat # 213477; Abcam), NGAL (Cat # MLCN20; R&D Systems), albumin (Cat # ab207620; Abcam), and creatinine (Cat # 1012; Ethos Biosciences) were measured using ELISA kits. Urine renin concentration was measured using a renin activity ELISA kit (Cat # IB59131; IBL-America) after incubating urine samples with additional recombinant mouse AGT.

### Two-photon intravital microscopy

Rat serum albumin (Millipore-Sigma, Burlington, MA) was conjugated to Texas Red-X-succinimidyl ester (Thermo Fisher Scientific, Waltham, MA) and two-photon intravital microscopy was performed as described previously.^16^ Detailed methods are described in the Supplemental Materials.

### Bulk RNA sequencing

RNA was extracted from mouse kidneys using Maxwell® RSC simplyRNA Tissue Kits (Cat # AS1340; Promega) in accordance with the manufacturer’s protocol. Total RNA samples were shipped to Novogene for bulk mRNA sequencing (n=6 biological replicates/group). cDNA library was generated from total mRNA (1 µg) using NEBNext UltraTM RNA Library Prep Kits for Illumina (New England BioLabs). cDNA libraries were sequenced by NovaSeq 6000 (Illumina) in a paired-end fashion to reach more than 1.5M reads. Paired-end read data formatted to FASTQ were mapped to mouse genome mm10 using STAR (v2.5, mismatch=2) and quantified using HTSeq (v0.6.1, -m union).^17,18^

### Statistical analysis

Data were presented as either mean ± SEM or median with the 25^th^ and 75^th^ percentiles depending on whether the data were analyzed by parametric or non-parametric tests. Normality and homogeneous variance assumptions for raw or log-transformed data with n>5/group were assessed using the Shapiro-Wilk test and the Brown-Forsythe test, respectively. Student’s t-test and one-way analysis of variance (ANOVA) with the Holm-Sidak post-hoc test were used for the data that met both normality and homoscedasticity to compare two-group and multiple-group (n≥3) means. Welch’s t-test was used for data that passed normality test, but failed to satisfy equal variance assumption to compare two-group means. For data that did not pass either normality or equal variance test, we applied Mann-Whitney U-test for two-group comparisons or Kruskal-Wallis one-way ANOVA followed by Dunn’s method for multiple-group comparisons. Albumin uptake described in Figure 7 was analyzed using a linear mixed effect model with unstructured covariance including genotype, time (10, 30, and 60 minutes), and interaction between genotype and time as main effects and intercept and time as random effects. SigmaPlot version 15 (SYSTAT Software Inc.) was used for statistical analysis except for the data presented in Figures 5 and 7. RNA sequencing data analysis in Figure 5 was performed using the edgeR Bioconductor package (v3.36.0) for differential gene expression (DEG) analysis and the clusterProfiler R Bioconductor package (v4.2.2) for gene ontology (GO) analysis. Data presented in Figure 7 were analyzed using the nlm R package (version 3.1) in R (version 4.2.2). Statistical significance was set at P<0.05.

## RESULTS

### Western diet feeding did not change renal megalin abundance in mice

We first determined whether Western diet feeding changes renal megalin abundance in mice. qPCR and Western blot analyses were performed to quantify megalin mRNA and protein abundance in the kidney of wild-type mice fed Western diet for 12 weeks (**Figure S2**). Renal megalin abundance was not changed by Western diet at either the mRNA or protein level.

### Validation of inducible PTC-specific megalin deletion in mice

To investigate the role of megalin in PTCs, we generated a mouse model using *Ndrg1-CreERT2*. NDRG1 protein has a predominant abundance in the renal cortex of PTCs (S1 and S2 segments).^9,19^. PTC-specific megalin deleted mice were generated using *Lrp2* floxed (*Lrp2* f/f) mice and *Cre* transgenic mice expressing a tamoxifen-inducible *Cre* recombinase under the control of *Ndrg1* promoter.^9,19,20^ Floxed mice in which exons 72-74 of *Lrp2* were flanked with *LoxP* sites (**Figure 1A**) were developed using the strategy reported by Leheste and colleagues ^8^. Male *Ndrg1*-*Cre ERT2*^+/0^ mice were bred with female *Lrp2* f/f mice to generate F1, F2 and littermates for *in vivo* studies (**Figure S1**) that had either of the two genotypes: *Ndrg1-Cre ERT2*^0/0^ *Lrp2* f/f (PTC-LRP2 +/+) or *Ndrg1*-Cre *ERT2*^+/0^ *Lrp2* f/f (PTC-LRP2 −/−). Offspring from F2 at 4-6 weeks of age were injected intraperitoneally with tamoxifen (150 mg/kg/day) for 5 consecutive days. Two or 15 weeks after completing the intraperitoneal injection of tamoxifen, cortex and medulla were isolated from kidney tissues to determine the floxed allele and deletion of megalin in the renal cortex (**Figure 1B**). qPCR confirmed significant reductions (~80%) of *Lrp2* mRNA in kidneys of PTC-LRP2 −/− mice (**Figure 1C**). As demonstrated by immunostaining for protein distribution, *Cre*-*LoxP* recombination led to deletion of megalin in S1 and S2 of PTCs, but its presence in S3 remained (**Figure 1D**).

**Figure 1.**
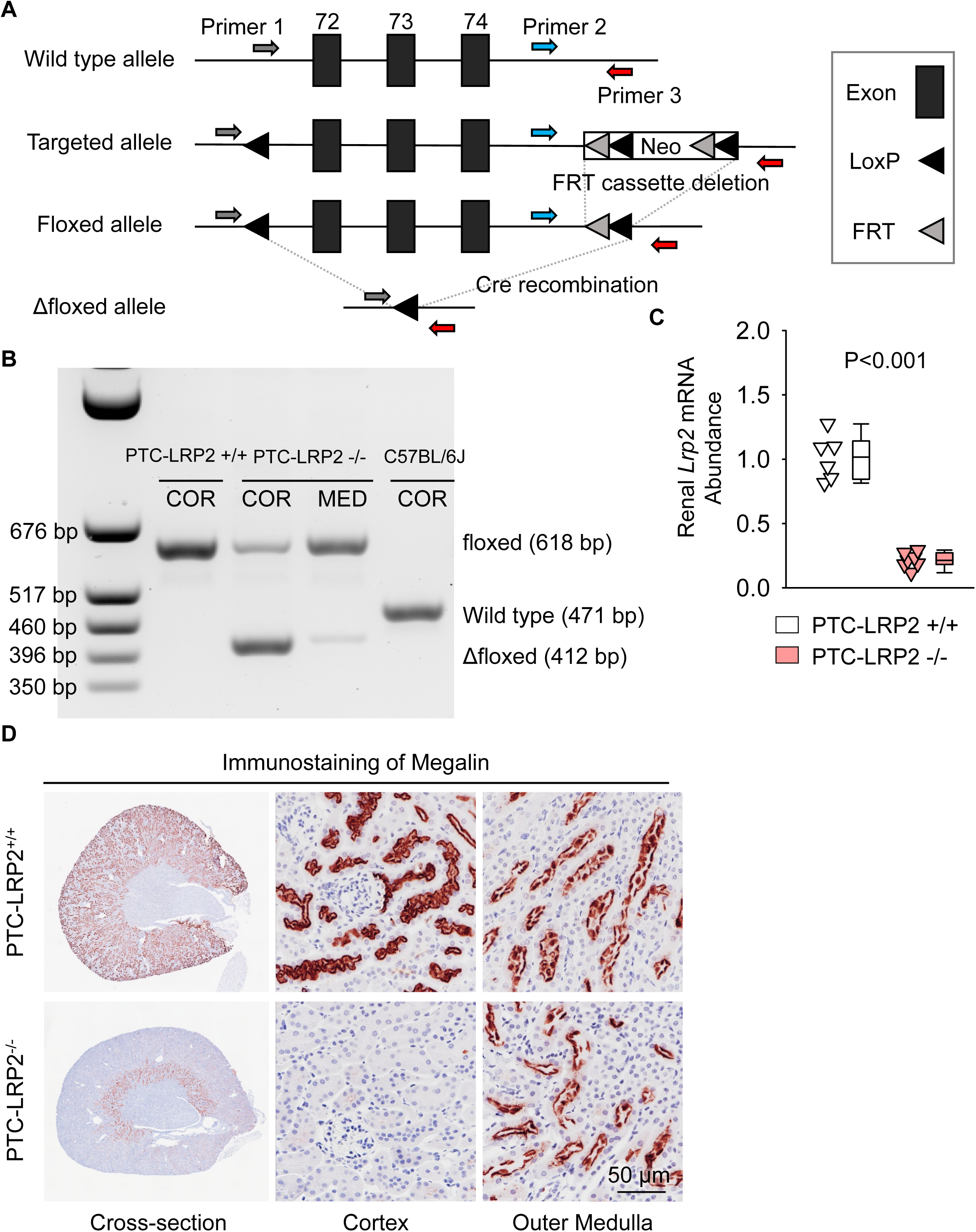
Development and validation of PTC-specific megalin deficient mice. **(A)** Construct map of the *Lrp2* floxed mouse. Three *Lox*P sites were inserted to encompass exons 72-74 of the mouse *Lrp2* gene. One *Lox*P site was inserted in intron 71, and 2 *Lox*P sites were in intron 74. A neo cassette in intron 74 was flanked by the 2 *Lox*P sites and 2 FRT sites in intron 74. After removal of the neo cassette, *Cre* recombination enabled the deletion of exons 72-74 of *Lrp2*, resulting in megalin deletion. **(B)** DNA-PCR using cortex (COR) and medulla (MED) regions of kidneys harvested from male wild-type (WT; C57BL/6J), *Ndrg1*-*Cre ERT2* 0/0 *Lrp2* f/f (PTC-LRP2 +/+), and *Ndrg1*-*Cre ERT2* +/0 *Lrp2* f/f (PTC-LRP2 −/−) mice 2 weeks after the completion of intraperitoneal tamoxifen injection. **(C)** Renal mRNA abundance of *Lrp2* was determined by qPCR (N=6-7/group), and analyzed using Welch’s t-test. **(D)** Immunostaining of megalin illustrated the distribution of megalin in kidneys of PTC-LRP2 +/+ and PTC-LRP2 −/− mice at 2 weeks after the completion of tamoxifen injections.

Our and others’ previous studies demonstrated that AGT, the substrate of all angiotensin peptides, in S1 and S2 of PTCs is derived primarily from hepatocytes, whereas AGT protein in S3 of PTCs is derived from kidney.^7,21^ In the absence of megalin in S1 and S2 of PTCs, AGT became abundant in S3 of PTCs, while its presence in S1 and S2 of PTCs was minimal (**Figure S3**).

Mass spectrometry-assisted proteomics analysis was performed on urine samples from male PTC-LRP2 +/+ and −/− mice at 2 weeks after completion of tamoxifen injection prior to Western diet feeding (**Figure S4**). The analysis identified 550 molecules, including 10 megalin ligands. While protein abundance of albumin (ALB), apolipoprotein E (APOE), and urokinase (PLAU) did not differ between the two genotypes, urinary excretion of multiple megalin ligands, such as AGT, retinol-binding protein 4 (RBP4), and lipoprotein lipase (LPL), lipocalin-2 (LCN2, also known as neutrophil gelatinase-associated lipocalin, NGAL), were elevated significantly in PTC-LRP2 −/− mice compared to their wild-type littermates. Of note, urinary REN1 was only detected in PTC-LRP2 −/− mice. These findings suggest a functional impairment of megalin as an endocytic receptor in PTC-LRP2 −/− mice.

### PTC-specific megalin deletion did not affect atherosclerosis in hypercholesterolemic mice

Following validation of the phenotype of genetically manipulated mice, we determined the effects of PTC-specific megalin deletion on blood pressure and atherosclerosis in both sexes (**Figure 2 and Figure S6**). PTC-LRP2 +/+ and PTC-LRP2 −/− littermates in an LDL receptor −/− background were injected with tamoxifen at 4-6 weeks of age. Two weeks after the completion of tamoxifen injections, Western diet feeding was started and continued for 12 weeks. Our previous study reported that *Lrp2* ASO reduced hypercholesterolemia-induced atherosclerosis in LDL receptor −/− mice.^7^ Therefore, subcutaneous injection of *Lrp2* ASO (6 mg/kg/week) to one group of PTC-LRP2 +/+ littermates was used as a positive control for this atherosclerosis study. Since our previous study has confirmed that control ASO showed comparable results as PBS (the solvent for ASOs) on blood pressure and atherosclerosis, PBS injection was used as the negative control (vehicle) of *Lrp2* ASO (**Figure 2A**). Western blot analyses using tissues from the kidney cortex demonstrated that megalin was modestly decreased by *Lrp2* ASO in PTC-LRP2 +/+ mice, whereas it was barely detected in PTC-LRP2 −/− mice (**Figure 2B**).

**Figure 2.**
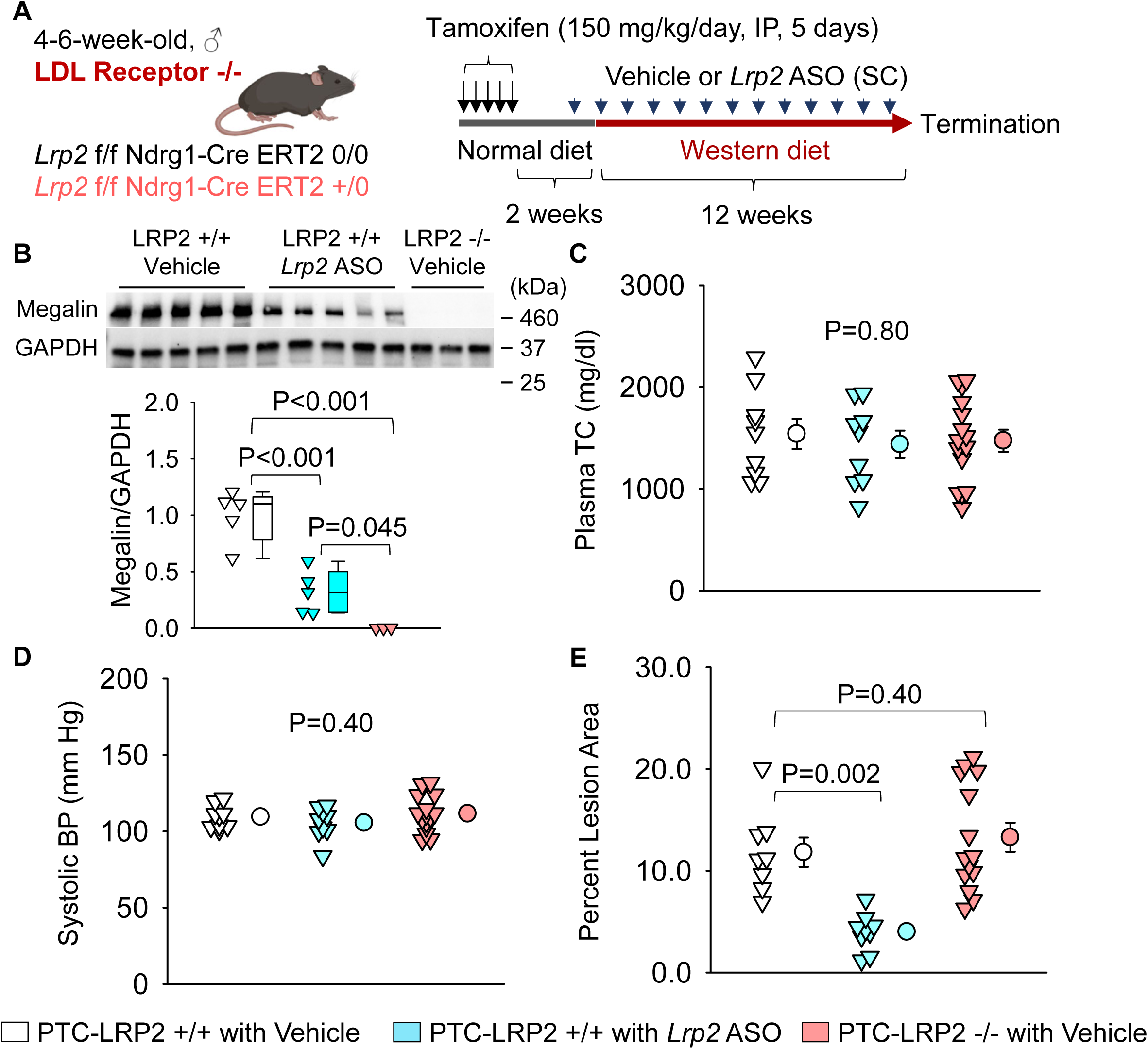
PTC-specific deletion of megalin did not attenuate hypercholesterolemia-induced atherosclerosis. **(A)** Experimental protocol: Four to 6-week-old male mice in an LDL receptor −/− background received intraperitoneal (IP) injections of tamoxifen for 5 consecutive days. Two weeks after completing the tamoxifen injection, all study mice were fed a Western diet for 12 weeks. The study mice received subcutaneous (SC) injections of PBS (Vehicle) or *Lrp2* antisense oligonucleotides (*Lrp2* ASO, 6 mg/kg/week) started 1 week prior to Western diet feeding. **(B)** Renal megalin protein abundance was determined by Western blot. **(C)** Plasma total cholesterol (TC) concentrations were measured using an enzymatic method. **(D)** Systolic blood pressures (BP) were measured using a tail-cuff system. **(E)** Percent atherosclerotic lesion area was quantified using an *en face* approach. Statistical analysis: Kruskal-Wallis one-way ANOVA on ranks followed by the Dunn method **(B)** and one-way ANOVA followed by the Holm-Sidak method **(C-E)**.

Immunostaining of megalin showed that megalin was reduced uniformly across all segments (S1, S2, and S3) of PTCs in mice with *Lrp2* ASO injection. In contrast, megalin was ablated in the S1 and S2 of PTCs, but remained in S3 of PTC-LRP2 −/− mice (**Figure S5**). Since megalin interacts with cubilin on PTCs, we determined whether cubilin is increased in a compensatory mode in response to megalin deletion. qPCR and Western blot analyses consistently demonstrated that cubilin was not enhanced in PTC-LRP2 +/+ mice administered *Lrp2* ASO or in PTC-LRP2 −/− mice (**Figure S6A, B**). Compared to PTC-LRP2 +/+ mice with vehicle administration, the distribution of cubilin in the kidney was not changed in PTC-LRP2 +/+ mice with *Lrp2* ASO injections or in PTC-LRP2 −/−mice (**Figure S7**).

Plasma total cholesterol concentrations were not different among the three groups in either sex (**Figure 2C, Figure S8A**). Also, inhibition by *Lrp2* ASO or PTC-specific deletion of megalin did not change systemic blood pressure (**Figure 2D, Figure S8B**). Consistent with our previous findings,^7^ inhibition of megalin globally by *Lrp2* ASO significantly suppressed atherosclerosis development in both male and female PTC-LRP2 +/+ mice (**Figure 2E, Figure S8C**). However, atherosclerotic lesion size was not different between PTC-LRP2 +/+ and PTC-LRP2 −/− littermates injected with PBS.

### PTC-specific megalin deficiency led to TIN in male LDL receptor −/− mice fed Western diet

Surprisingly, during necropsy, we noted that all male littermates with PTC-specific megalin deficiency exhibited smaller kidney weight and abnormal morphology with a distinctly irregular surface (**Figure 3A and 3B**). We did not observe overt morphological changes in male wild-type littermates injected with *Lrp2* ASO. Histological analysis was performed to illustrate the features of the renal pathologies in male PTC-LRP2 −/− mice fed a Western diet, compared to their PTC-LRP2 +/+ littermates. With hematoxylin and eosin staining (**Figure 3C**), the cortex and the outer medulla parts of kidneys obtained from male PTC-LRP2 +/+ mice were uniform, whereas weak staining of eosin presented in a radial pattern in the cortex of kidneys from male PTC-LRP2 −/− mice. This pattern was associated predominantly with PTC atrophy (**Figure 3C**). Additionally, there are many cells accumulated in the interstitial areas.

**Figure 3.**
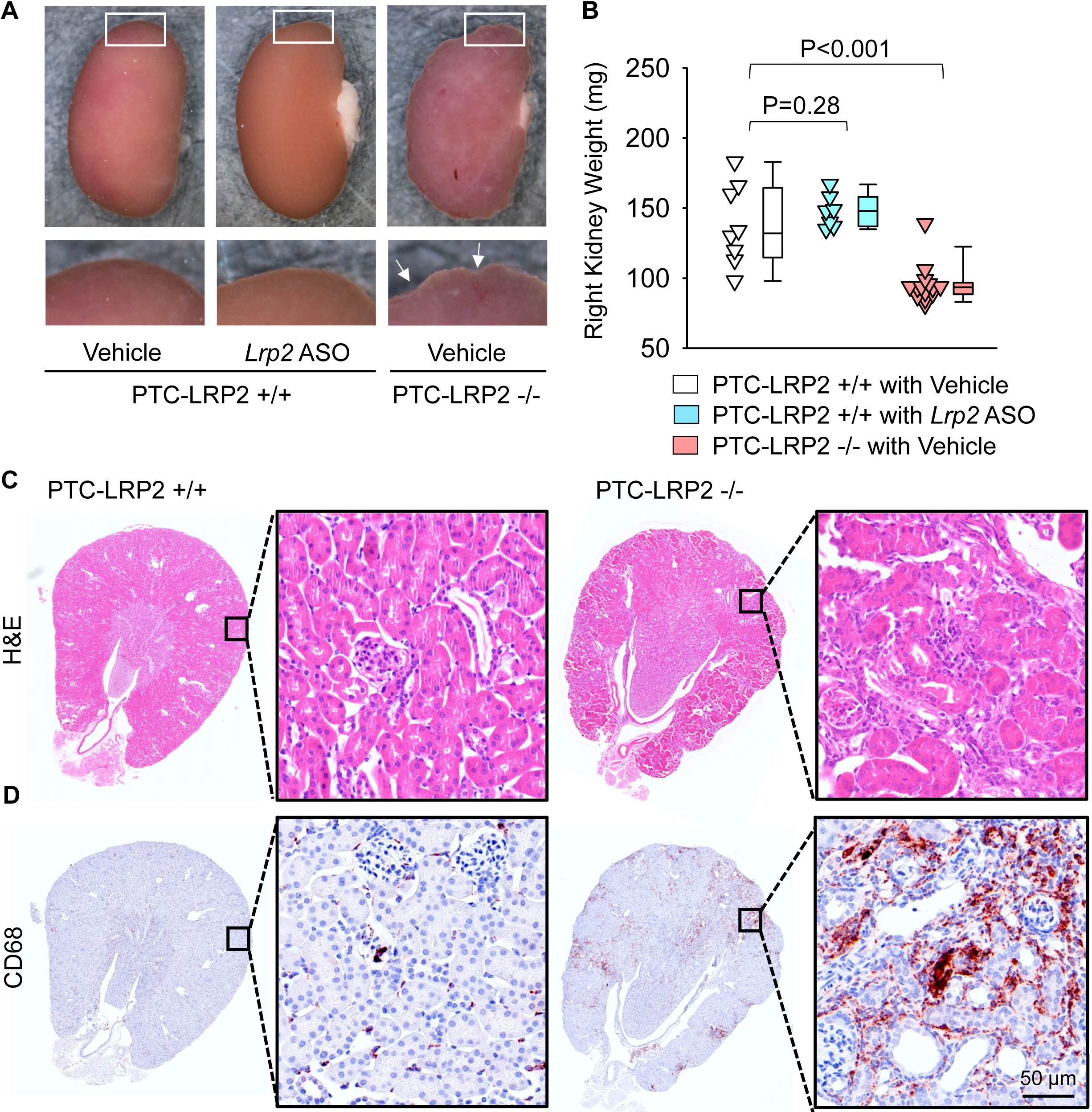
PTC-specific megalin deletion led to TIN in male LDL receptor −/− mice fed a Western diet. Four to 6-week-old male mice on an LDL receptor −/− background received intraperitoneal injections of tamoxifen for 5 consecutive days. Two weeks after completing the tamoxifen injection, all study mice were fed a Western diet for 12 weeks. The study mice received subcutaneous (SC) injections of PBS (Vehicle) or *Lrp2* antisense oligonucleotides (*Lrp2* ASO, 6 mg/kg/week) started 1 week prior to Western diet feeding. **(A)** Gross appearance of kidneys and **(B)** right kidney weight at termination. Statistical analyses: one-way ANOVA followed by the Holm-Sidak test after log-transformation. **(C)** Hematoxylin and eosin (H&E) staining, and **(D)** immunostaining of CD68 in kidney (N=7-14 per group).

Immunostaining of inflammatory cell markers revealed that CD45- and CD68-, but not CD3- or CD19-, positive inflammatory cells were accumulated in the cortex of PTC-LRP2 −/− mice after 12 weeks of Western diet feeding (**Figure 3D, S9**). To determine the impact of PTC-specific megalin deletion on other organs, immunostaining of CD68 was also performed in the heart and liver of PTC-specific megalin deficient mice and their wild-type littermates. There was no apparent macrophage accumulation in the heart regardless of genotypes (**Figure S10A**). Discernable differences of CD68 immunostaining were not apparent in the liver between genotypes (**Figure S10B**). These data suggest that PTC-specific megalin deletion leads to inflammation predominantly in kidney, but not in other organs. Of note, PTC-specific megalin deletion also led to renal fibrosis, which was spatially coincident with macrophage-accumulated areas (**Figure S11**). Despite the severe inflammation and fibrosis, significant glomerular damage was not observed in PTC-LRP2 −/− mice (**Figure S12**). Overall, these pathological features are consistent with TIN, a common cause of acute kidney injury that can progress to chronic kidney disease.^22^

In contrast to the severe pathological changes observed in male PTC-LRP2 −/− mice fed the Western diet, kidneys harvested from female PTC-LRP2 −/− mice after 12 weeks of the same diet did not exhibit morphological alterations. There were no discernable differences in kidney weight between the two genotypes (**Figure S13A, B**). H&E staining did not detect obvious proximal tubule atrophy in female PTC-LRP2 −/− mice (**Figure S13C**). Immunostaining revealed sparsely accumulated CD68+ cells in the interstitial space in both PTC-LRP2 +/+ and PTC-LRP2 −/− mice (**Figure S13D**).

To determine whether the renal pathologies were related to Western diet feeding, male mice with PTC-specific deletion of megalin were fed a normal laboratory diet for 15 weeks after completing injections of tamoxifen (**Figure S14A**). Plasma total cholesterol concentrations and kidney weight were not different between PTC-LRP2 +/+ and −/− mice (**Figure S14B, C**). Although deletion of megalin in S1 and S2 of PTCs was evident in male PTC-LRP2 −/− mice, no apparent renal pathologies were observed (**Figure S14D, E**).

In the absence of megalin in PTCs, AGT and renin were present in high concentrations in urine (**Figure 4A and B, Figure S15A and B**). RBP4, a functional biomarker of PTCs, is regulated by megalin.^7,23^ RBP4 in urine was not detectable in PTC-LRP2 +/+ mice; however, it was present in high concentrations in PTC-LRP2 −/− mice, irrespective of sex (**Figure 4C, Figure S15C**). Albumin, filtered through glomeruli, is normally taken up by renal PTCs in a megalin and cubilin-mediated manner.^24^ The ratio of urinary albumin to urine creatinine was increased by PTC-specific megalin deficiency in both male and female mice (**Figure 4D, Figure S15D**). Urine neutrophil gelatinase-associated lipocalin (NGAL) and kidney injury molecule-1 (KIM-1) are biomarkers representing impaired proximal tubules.^25^ Pronounced increases of urinary NGAL and KIM-1 were observed in both sexes of PTC-LRP2 −/− mice (**Figure 4E and F, Figure S15E and F**). Despite the loss of high concentrations of renin in urine, plasma renin concentrations did not differ among groups regardless of sex (**Figure S16**), suggesting that megalin deletion does not influence the systemic renin-angiotensin regulation. Overall, there were increased concentrations of many megalin ligands in the urine of PTC-LRP2 −/− mice.

**Figure 4.**
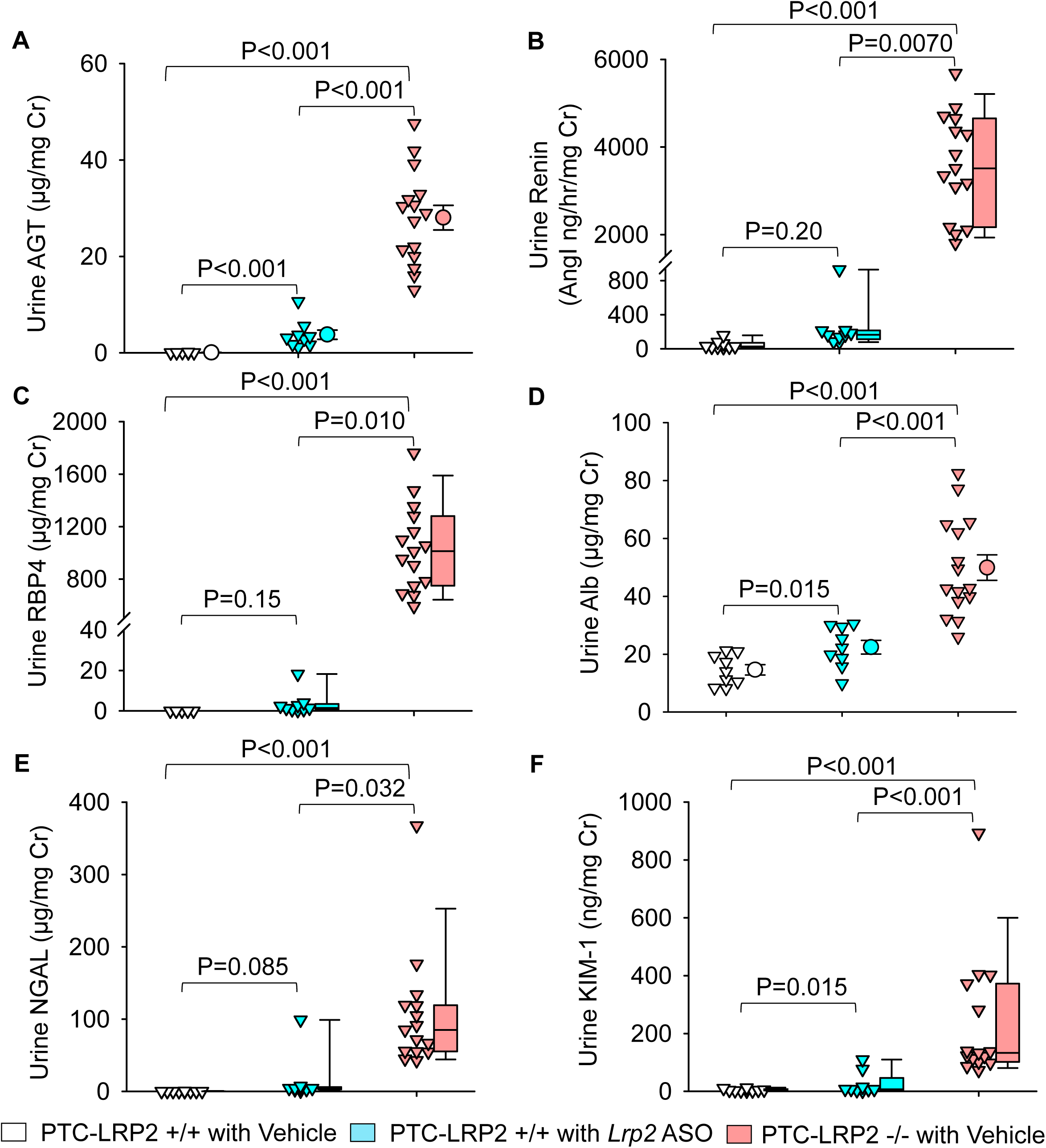
PTC-specific megalin deletion increased renal PTC injury markers in male LDL receptor −/− mice fed a Western diet. Four to 6-week-old male mice in an LDL receptor −/− background received intraperitoneal injections of tamoxifen for 5 consecutive days. Two weeks after completing the tamoxifen injection, all study mice were fed a Western diet for 12 weeks. The study mice received subcutaneous (SC) injections of PBS (Vehicle) or *Lrp2* antisense oligonucleotides (*Lrp2* ASO, 6 mg/kg/week) started 1 week prior to Western diet feeding. Urine was collected before termination. AGT **(A)**, renin **(B)**, RBP4 **(C)**, albumin **(D)**, NGAL **(E)**, and KIM-1 **(F)** in urine were measured using ELISA kits and normalized by urine creatinine concentrations. Statistical analysis: Mann-Whitney U-test **(A, C-F)** and Welch’s t-test **(B)** because data presented in **(B)** passed the normality but did not pass the equal variance test. Statistical analysis: one-way ANOVA with Holm-Sidak post hoc test **(A, D)** or Kruskal-Wallis one-way ANOVA on Ranks with Dunn post hoc test **(B, C, E, F)**.

### PTC-specific megalin deficiency augmented inflammation-related transcriptomes in kidneys of male LDL receptor −/− mice fed Western diet

To explore potential mechanisms by which deletion of PTC-specific megalin deletion induced renal pathologies, 2 weeks after tamoxifen induction, male PTC-LRP2 +/+ and PTC-LRP2 −/− littermates were fed a Western diet for either 2 weeks (representing early pathological status) or 12 weeks (representing advanced pathological status). Gross morphology was not apparently different between the two genotypes when the Western diet was fed only for 2 weeks. Renal cortex from each mouse was collected to isolate RNA and bulk RNA sequencing was performed subsequently. Transcriptomic patterns of the two genotypes at 2 weeks of Western diet feeding were different, as illustrated by principal component analysis (**Figure 5A**). PTC-specific megalin deletion resulted in up- and downregulation of 2959 and 2582 genes, respectively (**Figure 5B**). Enrichment analysis using the 2959 upregulated DEGs demonstrated that inflammation-related pathways were enriched predominantly **(Figure 5C)**. Major inflammatory molecules, such as *Il6, Lcn2, Il1b, Tnf, Clcl2,* and *Stat1,* were upregulated in the kidneys from PTC-LRP2 −/− mice (**Figure 5D**). In addition to inflammatory molecules, multiple collagen genes were upregulated in the kidneys from PTC-LRP2 −/− mice (**Figure 5D**). These inflammatory and collagen genes remained upregulated after 12 weeks of Western diet feeding (**Figure S17**). Based on the findings from both pathological and transcriptomic assessments at an early stage (2 weeks of Western diet feeding) and a chronic stage (12 weeks of Western diet feeding) of the renal phenotypes, inflammation and fibrosis occurred rapidly after starting Western diet feeding. This was also consistent with macrophage accumulation and collagen deposition observed in the interstitial space of kidneys from PTC-LRP2 −/− mice fed Western diet. Of note, these findings are consistent with a recent study in which megalin deletion in mouse kidney, driven by a *Cre* under the control of an *Emx*, resulted in renal inflammation and fibrosis when the mice were fed a similar Western diet.^26^

**Figure 5.**
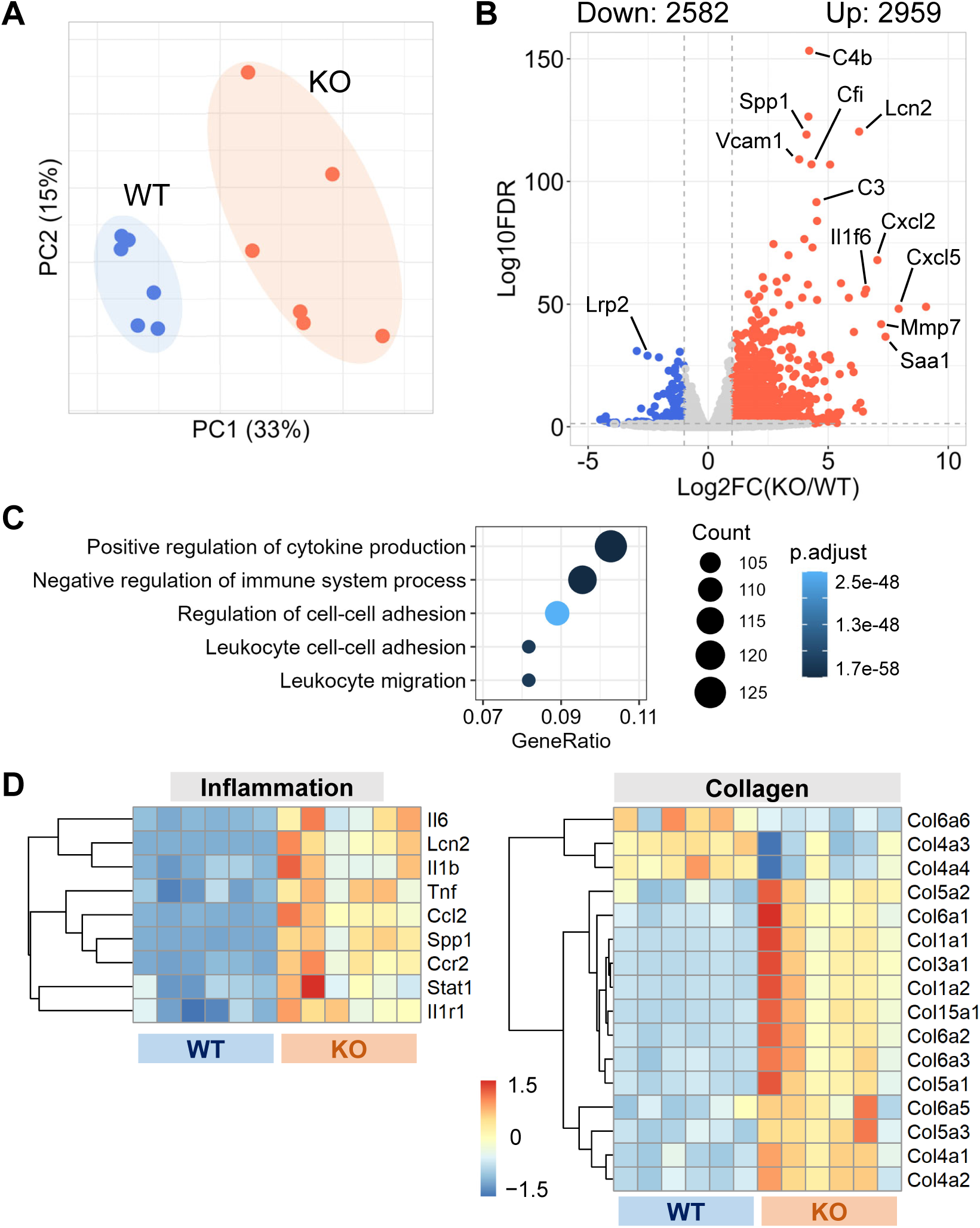
PTC-specific megalin deficiency augmented inflammation-related transcriptomes in male mice fed a Western diet. **(A)** Principal component analysis of transcriptomes of PTC-LRP2 +/+ (WT) and PTC-LRP2 −/− (KO) mice fed a Western diet for 2 weeks. **(B)** Volcano plot depicting differentially expressed genes (DEGs) between the 2 genotypes. **(C)** Gene ontology enrichment analysis for upregulated DEGs. **(D)** Heatmap with Z-scored coloring displaying DEGs associated with inflammation and collagens after 2 weeks of Western diet. N=6/group.

### PTC-specific megalin deficiency led to TIN in male C57BL/6J mice fed Western diet

Striking renal pathologies were observed in male PTC-LRP2 −/− mice with an LDL receptor −/− background fed the Western diet, but not in male mice fed a normal laboratory diet for comparable intervals. To determine whether the renal pathologies were dependent on LDL receptor deficiency, we repeated the study in LDL receptor +/+ mice that were on a strain-equivalent background of C57BL/6J (**Figure 6A**). PTC-LRP2 +/+ and PTC-LRP2 −/− littermates in LDL receptor +/+ background were injected with tamoxifen at 4-6 weeks of age. Two weeks after completion of tamoxifen injections, Western diet feeding was started and continued for 12 weeks. PTC-specific megalin deficiency in male LDL receptor +/+ mice showed abnormal renal morphology, proximal tubule atrophy, and interstitial inflammation (**Figure 6B-F**) that were consistent with the pathology observed in male PTC-LRP2 −/− mice in an LDL receptor −/− background, although these mice were not hypercholesterolemic. These data support the notion that Western diet feeding, rather than hypercholesterolemia, induces renal pathologies in male PTC-LRP2 −/− mice.

**Figure 6.**
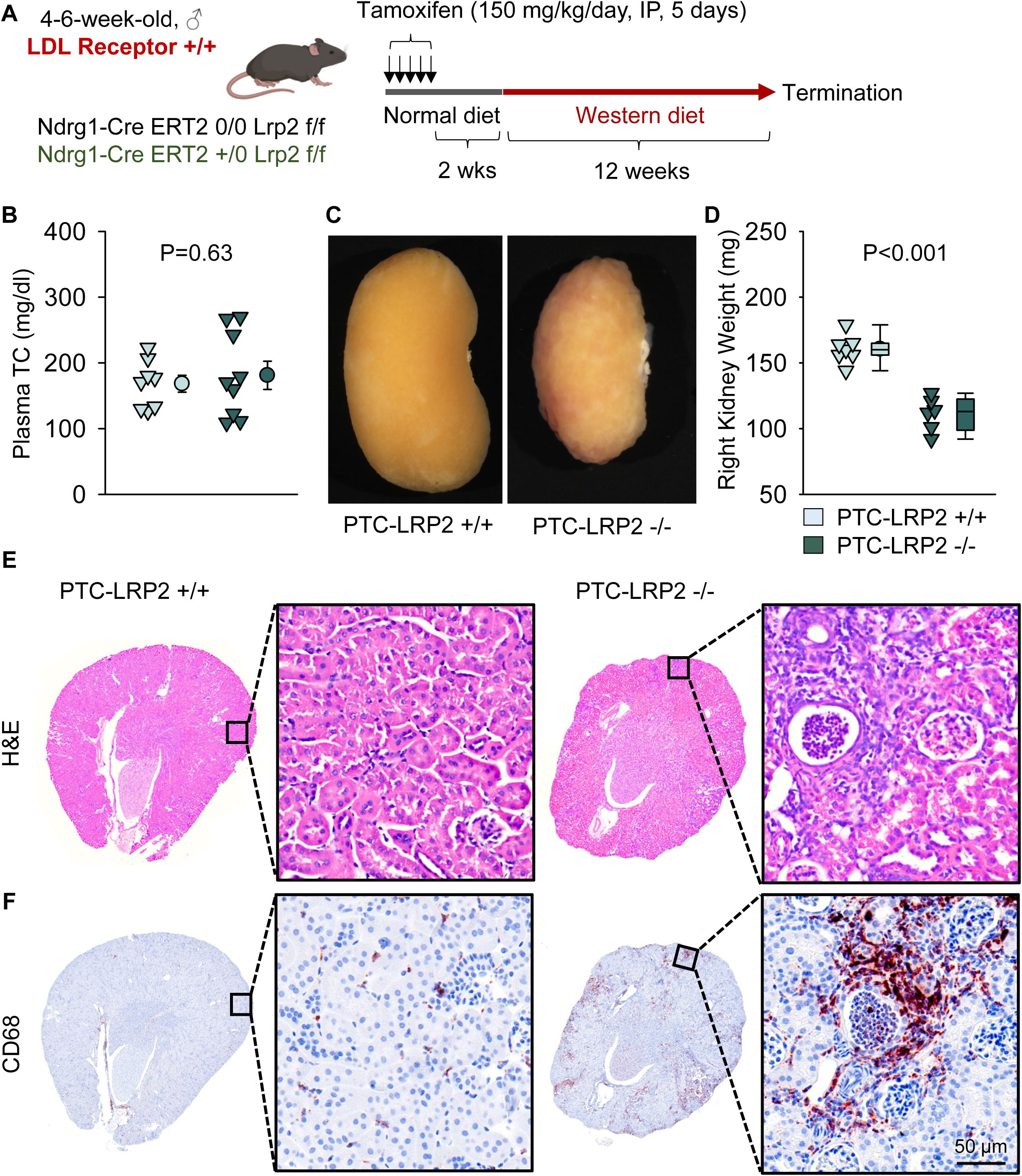
PTC-specific megalin deletion led to TIN in male LDL receptor +/+ mice fed a Western diet. **(A)** Experimental protocol: Five-week-old male LDL receptor +/+ mice on a C57BL/6J background received intraperitoneal (IP) injections of tamoxifen for 5 consecutive days. Two weeks after completing the tamoxifen injection, all mice were fed a Western diet for 12 weeks. **(B)** Plasma total cholesterol (TC) concentrations were measured using an enzymatic method. **(C)** Gross appearance of kidney, **(D)** weight of right kidney, **(E)** hematoxylin and eosin (H&E), and **(F)** immunostaining of CD68 in kidneys after termination. N=6-9/group. Statistical analysis: Student’s t-test **(B)** or Mann-Whitney Rank-Sum test **(D)**.

The renal pathologies, as determined by kidney weight, H&E staining, and immunostaining of CD68, were not apparent in female PTC-LRP2 −/− mice with the LDL receptor +/+ background fed Western diet (**Figure S18**).

### PTC-specific megalin deficiency led to impaired accumulation of fluorescently labeled albumin in male C57BL/6J mice fed a Western diet

Disruption of megalin function in PTC-LRP2 −/− mice in LDL receptor +/+ background was verified by quantitative intravital microscopy of proximal tubule uptake of albumin, a known ligand of megalin. After 10 days of Western diet feeding, male PTC-LRP2 +/+ and PTC-LRP2 −/− littermates had three-dimensional image volumes collected in vivo, before and at 10, 30, and 60 minutes, respectively, after intravenous injection of Texas Red-labeled rat serum albumin (TR-RSA). Representative fields collected from PTC-LRP2 +/+ and PTC-LPR2 −/− mice at each interval are shown in **Figure 7A**. Quantitative analyses demonstrated that the rate of initial uptake (measured at 10 minutes) was reduced ~47-fold in PTC-LPR2 −/− mice (**Figure 7B)**. Interestingly, significantly less punctate lysosomal autofluorescence (as shown by the pre-injection images in **Figure 7A**) was noted in PTC-LRP2 −/− mice, suggesting that loss of megalin might have decreased uptake of other endogenous ligands. In addition to the rapid disruption of albumin accumulation observed by intravital microscopy, 2 weeks of Western diet feeding also led to remarkable inflammation, as demonstrated by the accumulation of CD68 positive cells interstitially (**Figure S19**).

**Figure 7.**
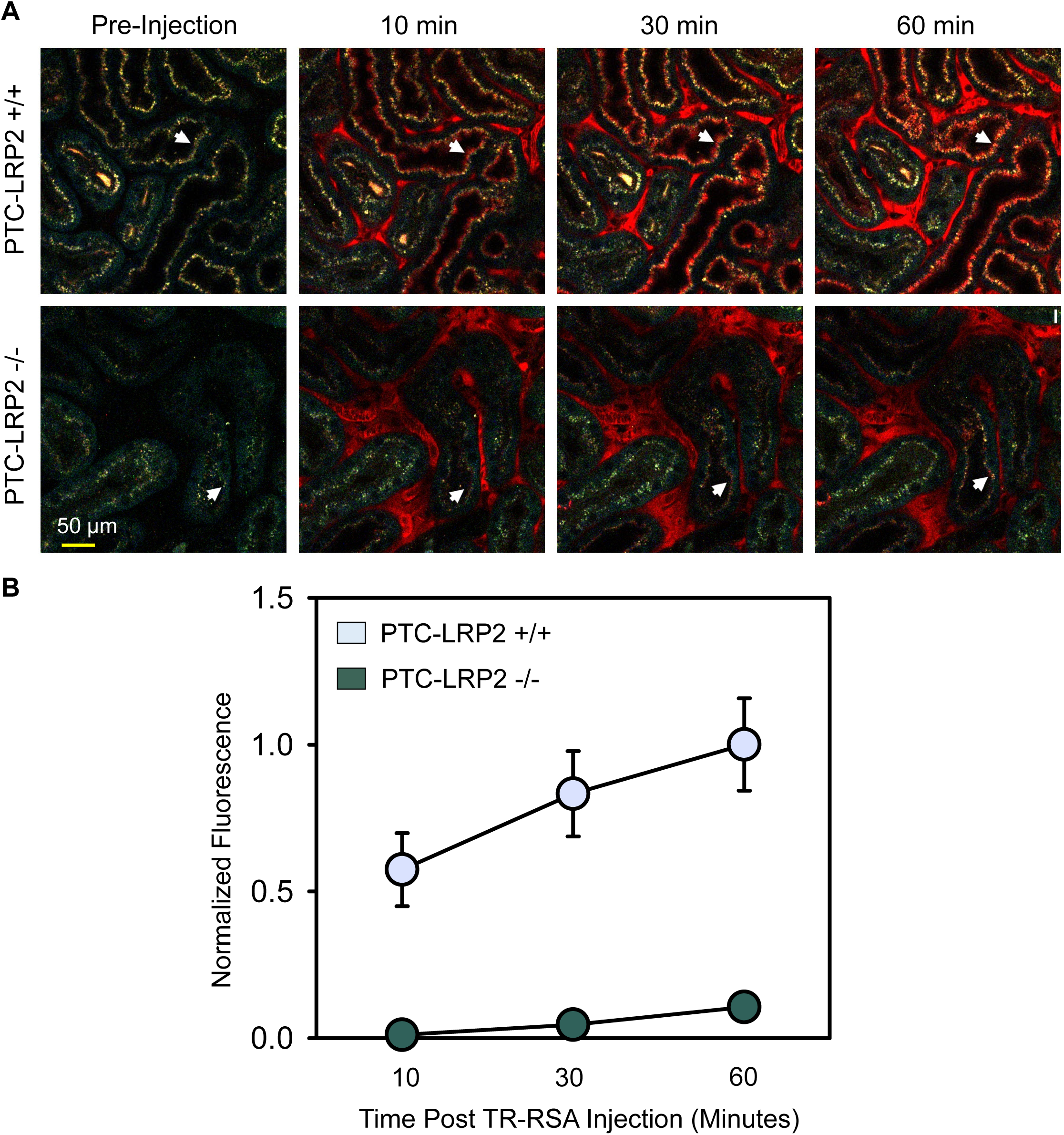
PTC-specific megalin deficiency led to impaired uptake of fluorescent albumin in male mice fed a Western diet. Four to 6-week-old male mice on an LDL receptor +/+ background received intraperitoneal injections of tamoxifen for 5 consecutive days. Two weeks after completing the tamoxifen injection, all study mice were fed a Western diet for 10 days. Multiphoton intravital microscopy was conducted to quantify PTC accumulation of albumin. **(A)** In vivo multiphoton fluorescence images were collected from the kidneys of PTC-LRP2 +/+ and PTC-LRP2 −/− mice prior to intravenous injection of Texas Red-labeled rat serum albumin (TR-RSA) (Top row), and 10, 30 and 60 minutes after injection. Arrows indicate examples of the same PTC regions imaged over time. **(B)** Quantitative analysis of albumin PTC uptake in images collected from PTC-LRP2 +/+ and PTCLRP2 −/− mice. Statistical analysis: a linear mixed effects model with unstructured covariance matrix. P=0.0034 between the two genotypes at 10 minutes, and P<0.001 for interaction between genotype and time.

## DISCUSSION

The primary objective of this study was to investigate the role of megalin in PTCs in hypercholesterolemia-induced atherosclerosis. Contrary to our initial hypothesis, deletion of megalin in S1 and S2 of PTCs did not reduce atherosclerosis in LDL receptor −/− mice. However, serendipitously, we observed that deletion of megalin in these two segments of PTCs led to TIN. There are several significant and novel findings shown in this study: (1) PTC-specific deletion of megalin resulted in TIN in male mice fed a Western diet, but not in mice fed a normal laboratory diet, (2) Western diet-induced TIN in PTC-LRP2 −/− mice was independent of hypercholesterolemia, (3) PTC-specific megalin deletion-induced TIN was severe in male mice, but not evident in female mice, and (4) PTC-specific megalin deletion resulted in rapid onset of interstitial inflammation and fibrosis following the initiation of Western diet feeding. This inflammation was accompanied by pronounced functional impairment of PTCs to uptake albumin, a prominent ligand of megalin.

Our previous study revealed that megalin is mainly present in renal PTCs of adult mice, while other tissues and organs either lack or have a very low abundance of megalin.^7^ Megalin is necessary to retain AGT and renin in renal PTCs, where high concentrations of AngII, a major contributor to blood pressure regulation and atherosclerosis, are present.^7,27–29^ Our previous and the present studies demonstrated consistently that ASO-induced deletion of megalin reduced atherosclerosis.^7^ Therefore, it was initially anticipated that PTC-derived megalin would be the primary source contributing to atherosclerosis. However, the present study involving a large number of animals, including both males and females, does not support this initial hypothesis. Effects of tamoxifen on atherosclerosis have been implicated in previous reports.^30^ Since all study mice were administered an equivalent amount of tamoxifen, the present study does not support the notion that transient tamoxifen administration affected atherosclerosis.

The basis for the lack of protection against atherosclerosis development following PTC-specific deletion of megalin is not entirely clear. Several differences between *Lrp2* ASO administration and PTC-specific megalin deficiency may contribute to these conflicting findings. The protein abundance reduction of megalin differed between the ASO pharmacological approach and the genetic deletion approach. Specifically, megalin protein abundance across S1, S2, and S3 was modestly reduced by *Lrp2* ASO, whereas megalin protein in S1 and S2 was abolished in genetically engineered PTC-LRP2 −/− mice. *Lrp2* ASO may preserve a small but crucial amount of megalin in S1 and S2 segments, unlike the complete deletion in PTC-LRP2 −/− mice. In addition, megalin was abundant in S3 following its genetic deletion in S1 and S2 of PTCs. It is unclear whether ablation of megalin in S1 and S2 is detrimental, or the high abundance of megalin in S3 plays a critical role in contributing to atherosclerosis. In a preliminary study, we attempted to delete megalin specifically in the S3 segment using *Kap2*-Cre, but failed to demonstrate a reduction. Unfortunately, we are not aware of other Cre promoters that target S3 specifically. Therefore, it is not technically feasible to delete megalin specifically in S3 of the kidney. Second, although loss of AGT, renin, RBP4, and albumin were also detected in mice injected with *Lrp2* ASO, no severe renal dysfunction was observed in these mice. In contrast, genetic deletion of megalin in S1 and S2 of PTCs in male mice led to severe renal impairment when fed Western diet. Since persistent kidney damage is an independent risk factor for atherosclerosis,^31–33^ it is possible that chronic renal dysfunction could contribute to atherosclerosis development in PTC-LRP2 −/− mice. Third, despite the low abundance, other organs, such as the lung and parathyroid gland also express megalin.^7^ Given that ASO affects systemically, we cannot rule out that other organs expressing megalin do not influence the development of atherosclerosis.

The most surprising finding in this study is that PTC-specific megalin deletion resulted in TIN in male mice fed a Western diet, but not when fed a normal laboratory diet. This striking phenotype was observed in both LDL receptor +/+ and LDL receptor −/− mice that are on a C57BL/6J background. Plasma total cholesterol concentrations in LDL receptor −/− mice fed a Western diet were greater than 1,000 mg/dl, but were less than 200 mg/dl in LDL receptor +/+ mice fed a Western diet. These results support the concept that currently unidentified constituents of Western diet are contributing to PTC-specific megalin deficiency-induced TIN, whereas hypercholesterolemia *per se* is not essential. The RNA sequencing analyses conducted on kidney samples from mice fed a Western diet for 2 or 12 weeks revealed notable differences between male PTC-LRP2 +/+ and PTC-LRP2 −/− mice, particularly evident in increased gene expression associated with inflammation and fibrosis. Increased inflammatory gene expression was associated with a pronounced accumulation of immunostained macrophages in the renal interstitial regions of PTC-LRP2 −/− mice. This finding starkly contrasts with the sparse presence of macrophages in the kidney of PTC-LRP2 +/+ mice, as well as in mice of either genotype fed a normal laboratory diet. Inflammation is recognized as a hallmark of TIN.^22^ In the present study, the early onset of inflammation, as evidenced after just 2 weeks of Western diet feeding in PTC-LRP2 −/− mice, indicates that inflammation may be a potential causal factor contributing to the renal pathologies observed in these mice. However, the precise mechanisms triggering TIN remain unclear.

The findings of renal pathologies in PTC-LRP2 −/− mice fed Western diet conflict with those reported by Kuwahara and colleagues that kidney-specific reductions of megalin improved high-fat diet-induced renal pathological impairment.^34^ In that study, *Lrp2* floxed mice were bred with transgenic mice expressing *Cre* under the control of a human *Apoe* promoter which led to megalin deletion in a mosaic pattern in mouse renal cortex, with ~50-60% of megalin remaining in most PTCs.^8,21,27,34^ C57BL/6J mice fed a high-fat diet (60% calories/wt from fat) for 12 weeks resulted in modest hypertrophy, lipid peroxidation, and cellular senescence of PTCs. These renal changes were attenuated with the mosaic deletion of megalin. In addition to the different diet contents, one major difference between our study and the previously reported studies is the magnitude of megalin deletion in PTCs. *Cre* transgene driven by a *Ndrg1* promoter led to ablation of megalin in S1 and S2 of PTCs, but *Apoe*-*Cre* transgene led to only ~40-50% deletion of megalin in most PTCs. Megalin mediates endocytosis of a variety of molecules in PTCs such as RBP4 and albumin; however, it may also mediate the uptake of harmful components in the saturated fat-enriched diet that should be eliminated to maintain physiological homeostasis. The renal pathologies observed in PTC-LRP2 −/− mice fed the Western diet were not detected in C57BL/6J mice administered *Lrp2* ASO, which inhibits megalin globally, but did not abolish megalin in S1 and S2 of PTCs. Megalin is abundant in S3 of PTCs in PTC-LRP2 −/− mice. S3 is sensitive to stimulations such as hypoxia.^35,36^ It is well established that defective fatty acid oxidation of PTCs in S3 augments inflammation, cell death, and fibrosis. The present study did not address the question of whether TIN pathology was directly attributable to the absence of megalin in S1 and S2 or the highly abundant megalin in S3. In addition to development of mice expressing Cre specifically targeting S3, single nucleus RNA-sequencing in combination with spatial transcriptomics would provide valuable insights into resolving this dilemma.

In addition to the renal pathologies observed in this study, adenine diet-induced TIN in mice or rats has been used frequently to study mechanisms and potential therapeutic strategies of TIN.^37,38^ In this model, adenine in the diet leads to obstruction of the urinary tract due to the precipitation of adenine crystals. Consequently, the obstructed urinary tract results in injury to renal tubules including proximal tubules.^37,39^. This model presents several limitations, including the significant variability in the severity of pathologies, which challenges its consistency in studying TIN. In contrast, the renal pathologies observed in the present study were remarkably consistent, occurring in all male mice with PTC-specific megalin deletion fed a Western diet – a diet that represents the current diet habits in many Western countries. Notably, the striking renal pathologies observed in PTC-LRP2 −/− mice mirrored kidney pathologies found in various cardiovascular diseases, such as hypertension and diabetes.^40^ However, no preclinical and clinical studies have reported whether megalin impairment in PTCs manifests under these prevalent cardiovascular disease conditions. Additionally, PTC-specific megalin deletion in mice represents some relatively rare immune complex conditions found in humans. Recent human case reports or observational studies have identified manifestations of TIN related to autoantibodies against megalin on the PTC brush borders, a condition termed “anti-megalin nephropathy”.^41–46^. This nephropathy presents as severe tubulointerstitial injury and inflammation, with no or mild glomerular impairment, yet ultimately progresses to advanced renal disease. Although the potential connection between fat-enriched diet feeding in PTC-LRP2 −/− mice and this human condition is unclear, megalin impairment-induced TIN in mice has potential relevance to humans, highlighting its importance in exploring potential molecular mechanisms underlying human diseases.

Although urinary protein concentrations were similarly severe between male and female mice, the pathological changes were not evident in female PTC-LRP2 −/− mice. This sex difference is also noted in the mouse model with adenine-induced TIN.^47^ It is well known that cardiovascular diseases have strong sex differences.^10^ Some human studies also suggest that renal dysfunction progresses more slowly in women than in men, although conflict findings exist.^48–52^. Sex hormones, such as estrogen, are potential contributors to this sexual dimorphism.^53,54^ A recent study revealed sexual dimorphism of transcriptomic alterations in response to androgen receptor activation in proximal tubules using multiple omics approaches.^55^ However, their impact on the development and pathogenesis of TIN has not been defined. Future studies will be needed to explore the potential mechanisms by which PTC-specific megalin deletion in male mice leads to more severe TIN.

In summary, deletion of megalin specifically in S1 and S2 of PTCs failed to mitigate hypercholesterolemia-induced atherosclerosis, but instead induced TIN with severe pathological changes in male mice. The consumption of a Western diet exerted a crucial role in triggering the observed TIN. Future studies aim to understand the potential molecular mechanisms and pathogenesis of TIN associated with megalin deletion in S1 and S2 of PTCs, related sexual dimorphism, as well as its long-term impact on kidney and cardiovascular functions.

## Abbreviations

AngII: angiotensin II
ASO: antisense oligonucleotides
LDL: low-density lipoprotein
LRP2: low-density lipoprotein receptor-related protein 2
PTC: proximal tubule cell
TIN: Tubulointerstitial nephritis

## Acknowledgments

Histological and immunohistochemical images were acquired using Zeiss Axioscan Z1 or 7 in the Light Microscopy Core at the University of Kentucky.

## Supplemental Material

Major Resources Tables

Online Figures S1-S19

## HIGHLIGHTS

1. Deletion of megalin specifically in S1 and S2 of PTCs (proximal tubules) does not reduce atherosclerosis in hypercholesterolemic mice, irrespective of sex.
2. Deletion of megalin in S1 and S2 of PTCs induces TIN (tubulointerstitial nephritis) with severe renal pathological changes in male mice.
3. PTC-specific megalin deficiency-induced TIN occurs in both male LDL receptor −/− and LDL receptor +/+ mice fed a Western diet.
4. The consumption of a Western diet exerts a crucial role in triggering the observed TIN in male mice with PTC-specific megalin deletion.

