## Supplemental Materials for "Renal Proximal Tubule Cell-specific Megalin Deletion Does Not Affect Atherosclerosis But Induces Tubulointerstitial Nephritis in Mice Fed Western Diet"

*These authors contributed equally

***SUPPLEMENTAL METHODS***

***Urinary Proteomics***

Trichloroacetic acid precipitation was performed to concentrate proteins and remove contaminants from urine samples. Briefly, trichloroacetic acid (15 µl) was added to 150 µl of samples, vortexed, and incubated at −20°C for 30 minutes. Samples were then centrifuged, and the supernatant was removed. The pellet was washed with 500 µl of ice-cold acetone and incubated at −20°C overnight. Subsequently, samples were centrifuged, and the supernatant was removed. Pellets were air-dried and resuspended in lysis buffer.

Protein concentrations were determined using the Cytiva 2-D Quant kit according to the manufacturer's protocol. A standard curve was prepared using Bovine serum albumin. Sample tubes were prepared containing 1 µl or 10 µl of sample. Precipitant and co-precipitant solutions were added to each tube, vortexed, and centrifuged. The supernatant was removed. Copper solution, de-ionized water, and working color reagent were added, and samples were incubated at room temperature for 15 minutes. The absorbances were read at 480 nm and protein concentrations were determined by generating a standard curve and plotting the absorbance of the standards against the quantity of the protein.

Mouse urine samples were digested enzymatically using the S-Trap™ micro-MS sample prep kit: ≤ 100 µg according to the manufacturer's protocol. In brief, equivalent protein amounts (20 µg) were digested for each sample. Proteins were diluted to yield a final volume of 23 µl with lysis buffer. Reduction solution was added, and samples were incubated at 55°C for 15 minutes. Alkylation solution was added, and samples were incubated at room temperature for 10 minutes. Following incubation, protein samples were added to S-trap micro columns provided in the kit to trap and desalt the proteins according to the manufacturer's instructions. For digestion, trypsin stock solution was diluted in TEAB (50 mM) at 1 mg/ml, and 1 µg of trypsin solution combined with 20 µl of 50 mM TEAB digestion buffer was added to the S-Trap microcolumn and proteins were incubated at 47°C for 2 hours to facilitate enzymatic digestion.

Following digestion, peptides were eluted from the S-Trap columns and dried to completion by vacuum centrifugation. Peptides were resuspended in acetonitrile (5% vol/vol), formic acid (0.1% vol/vol), and water prior to LC-MS/MS analysis.

Peptides were analyzed on a Thermo Scientific™ Orbitrap Exploris 240 mass spectrometer with an EASY-Spray source housing and a Vanquish Neo UHPLC system (Thermo Scientific™). Peptides were separated on an EASY-Spray HPLC column (150 mm x 75 µm, 2 µm particle size) paired with a PepMap™ Neo 5 µm C18 300 µm x 5 mm Trap Cartridge. Mobile phase A consisted of formic acid (0.1% vol/vol) in water and mobile phase B consisted of formic acid (0.1% vol/vol) in acetonitrile. The flow rate was set to 0.3 µl/min. Peptides were separated over a 90-minute linear gradient from 2-55% mobile phase B. The column was equilibrated for 1.5 minutes at mobile phase B (2%). Mobile phase B was increased to 10% over 10 minutes and increased again to 25% over the next 35 minutes. Mobile phase B was increased to 35% over 25 minutes followed by 55% mobile phase B for 15 minutes. Following peptide separation, the column was washed at 0.75 µl/min at 98% mobile phase B for 8 minutes, followed by column re-equilibration at 2% mobile phase B for 5 minutes.

Data was collected using a data-dependent acquisition (DDA) strategy. The instrument was set to 60,000 resolution, with top N precursor ions in a 3-second cycle time. Data was collected using positive ionization including charge states 2-4. Full scan (MS1) settings were set as follows: (a) scan range: 375-1200 m/z, (b) RF lens (%): 45, (c) AGC target: custom, (d) normalized AGC target (%) 250, (e) maximum injection time mode: custom, and (f) maximum injection time (ms): 20. Peptide fragmentation spectra (MS2) were collected using a normalized HCD collision energy set to 26%, 2 m/z isolation window, 15,000 resolution, normalized AGC target set to 50%, and automatic maximum injection time.

Data were analyzed against the Mus musculus UniProt database (downloaded March 2024) using the Thermo Scientific™ Proteome Discoverer (version 3.1) software and the SEQUEST search algorithm. The precursor mass tolerance was set to 10 ppm and a 0.6 Da fragment mass tolerance. Dynamic modifications included N-terminal acetylation (+42.011), N-terminal methionine oxidation (131.040), and methionine oxidation (+15.995). Carbamidomethylation (+57.021 Da) of cysteine amino acids was included as a fixed modification. The feature mapper node was enabled and performed retention time alignment with a maximum RT shift of 10 minutes and minimum S/N threshold set to 5. Precursor peptide abundances were based on intensities. Unique and razor peptides were used for quantification and only proteins assigned to a protein group were used for quantification. Normalization was performed based on the total peptide amount.

#### *In vivo uptake of fluorescent albumin in mouse kidneys*

##### Conjugation of fluorescent albumin

Rat serum albumin (Millipore-Sigma, Burlington, MA) was conjugated to Texas Red-X-succinimidyl ester (Thermo Fisher Scientific, Waltham, MA). Briefly, conjugates were prepared per the manufacturer's instructions and then dialyzed extensively against 5 x 4 L changes of saline (0.9% wt/vol) over 2 days at 4°C. Purified Texas Red labeled albumin (TR-RSA) was aliquoted into 10 mg tubes, lyophilized, and stored at −80 °C.

##### Mouse preparation/surgery for intravital microscopy

Mice were placed in an induction chamber connected to an anesthesia circuit, dispensing isoflurane at 2 to 4% (vol/vol) at a flow rate of 1 L/min $O_2$. Once the mouse was stabilized, the left side of the body above the kidney and the right side of the neck was shaved and cleaned. A 1 cm right ventral incision was made in the neck, the jugular was exposed, and all fat and fascia surrounding were cleared. The anterior end of the jugular was tied using a 4-0 silk suture to prevent bleeding. A small nick was made in the jugular vein, and a catheter was slid roughly 1 cm into the jugular vein and secured at the posterior end using a 4-0 suture. The catheter was sutured and secured to the skin in three different places. The renal surface was imaged by first making a small incision above the kidney and exteriorizing the kidney gently by gripping the fat from the lower pole and gently pulling out while squeezing the incision behind the kidney. The mouse was transferred over and placed on a second anesthesia circuit delivering isoflurane (2% vol/vol). A two $mm^2$ piece of gauze soaked in saline was used to stabilize the kidney in the center of a 50 mm diameter coverslip bottom dish with a 40 mm diameter coverslip (Willco Wells, Electron Microscopy Sciences, Hatfield, PA). Once centered and stabilized, a rectal probe was placed to monitor body temperature that was kept between 36 and 37°C. A lightweight black plastic cloth was placed over the mouse and a 2.5 $cm^2$ space was placed around the mouse, being used to support the heating pad set at medium placed over the mouse.

**Two-photon intravital microscopy**

Intravital imaging studies of the renal surface were conducted using a Leica Dive SP-8 (Leica Microsystems, Wetzlar, Germany), with a 40x water immersion objective (NA 1.1). Two-photon excitation at 800 nm was accomplished using a Mai-Tai mode locked laser. Blue, green, and red emissions were collected by the system onto separate 12-bit detectors. Although Texas Red was the only fluorophore utilized in the study, the other channels were collected to acquire a multi-color image of the mouse kidney which includes its autofluorescent signature. To assess accumulation of filtered TR-RSA by proximal tubules, 8 regions containing mostly proximal tubules were marked and background images were collected for each region. A separate region with prominent vessels was selected and ~0.5 mg of TR-RSA was slowly infused while acquiring a time series to assure the fluorescence in the plasma is kept just below saturation. Subsequent images for each region were acquired 10, 30, and 60 minutes after injection of TR-RSA for analysis.

**Image analyses**

TR-RSA accumulation was quantified in images of 8 tubular image fields collected at each time point from the kidneys of PTC-LRP2 +/+ and -/- mice. In each field, ~9 tubular regions were carefully outlined and the mean intensity in the TR-RSA channel was measured using Metamorph image processing software (San Jose, CA). For each time point, TR-RSA fluorescence of each region was quantified as the mean intensity less the background fluorescence of that same region, as measured in the mean intensity measured in the image collected prior to TR-RSA injection. The analyzed data (a total of ~288 tubules from each) were normalized to the highest value (obtained from the PTC-LRP2 +/+ group at 60 minutes)

### MAJOR RESOURCES TABLES

#### Table I. Breeding strategy for *Ndrg1-Cre ERT2 Lrp2* f/f mice

| | Male parent | Female parent | Offsprings |
|---|---|---|---|
| F0 | *Ndrg1-Cre ERT2* +/0 | *Lrp2* f/f | Female f/+ x *Ndrg1-Cre ERT2* 0/0<br>Male f/+ x *Ndrg1-Cre ERT2* +/0 |
| F1 | *Ndrg1-Cre ERT2* +/0<br>*Lrp2* f/+ | *Ndrg1-Cre ERT2* 0/0<br>*Lrp2* f/+ | Female f/f x *Ndrg1-Cre ERT2* 0/0<br>Male f/f x *Ndrg1-Cre ERT2* +/0 |
| F2 | *Ndrg1-Cre ERT2* +/0<br>*Lrp2* f/f | *Ndrg1-Cre ERT2* +/0<br>*Lrp2* f/f | f/f x *Ndrg1-Cre ERT2* 0/0<br>f/f x *Ndrg1-Cre ERT2* +/0<br>Both males and females were used for experiments. |

#### Table II. Animals (in vivo studies) – In house breeding (Littermates)

| Mouse Strain | Background Strain | Sex |
|---|---|---|
| *Ndrg1-Cre ERT2* 0/0 *Lrp2* f/f (PTC-LRP2 +/+) | LDL Receptor -/- | Male & Female |
| *Ndrg1-Cre ERT2* +/0 *Lrp2* f/f (PTC-LRP2 -/-) | LDL Receptor -/- | Male & Female |
| *Ndrg1-Cre ERT2* 0/0 *Lrp2* f/f (PTC-LRP2 +/+) | C57BL/6J | Male & Female |
| *Ndrg1-Cre ERT2* +/0 *Lrp2* f/f (PTC-LRP2 -/-) | C57BL/6J | Male & Female |

#### Table III. Primers for qPCR

| Gene | Forward Primer | Reverse Primer |
|---|---|---|
| *Actb* | GCCTTCCTTCTTGGGTATGG | GCACTGTGTTGGCATAGAGG |
| *Gapdh* | CAACTCCCACTCTTCCACCT | CTTGCTCAGTGTCCTTGCTG |
| *Rplp2* | ATGTCATCGCTCAGGGTGTT | CTCCTCGGACTCCTCCTTCT |
| *Lrp2* | AGAATGTGGCAGTGGGAATTT | GGACAGCCAATTTCATCAGTGT |
| *Cubn* | TGTGAACTGTTTCTGGGTTGTC | TCTACCAAGTGGGAAATCTGC |

#### Table IV. Diets from Inotiv used in mouse studies

| Diet # | 2918 (Normal Laboratory Diet) | TD.88137 (Western Diet) |
|---|---|---|
| Carbohydrate (kcal/kcal%) | 58 | 42.7 |
| Protein (kcal/kcal%) | 24 | 15.2 |
| Fat (kcal/kcal%) | 18 | 42 |

#### Table V. Primary antibodies for immunostaining

| Antibody | Vendor | Cat # | Working Concentration |
|---|---|---|---|
| Rabbit anti-mouse/rat angiotensinogen (AGT) | IBL-America | 28101 | 1 μg/mL |
| Rabbit anti-CD68 (E307V) | Cell Signaling Technology | 97778 | 0.1 μg/mL |
| Rabbit anti-Megalin | abcam | ab76969 | 0.5 μg/mL for immunostaining, 0.1 μg/mL for Western blot |
| Rabbit anti-Cubilin | Invitrogen | PA5-115063 | 1 μg/mL for Western blot |
| Rabbit anti-GAPDH | Cell Signaling | 2118 | 0.1 μg/mL |

| | | | |
|---|---|---|---|
| (14C10) | Technology | | |
| Rabbit anti-CD3 | abcam | ab16669 | 1 µg/mL |
| Rabbit anti-CD19 | Cell Signaling Technology | 90176S | 1 µg/mL |
| Rat anti-CD45 | BD Pharmingen | 553076 | 0.3 µg/mL |

**Secondary Antibodies for immunostaining**

| **Antibody** | **Vendor** | **Cat #** |
|---|---|---|
| ImmPRESS goat anti-rabbit IgG | Vector | MP-7451 |
| ImmPRESS goat anti-rat IgG (Mouse Absorbed) | Vector | MP-7444 |
| Goat Anti-Rabbit IgG Antibody (H+L) | Vector Laboratories | PI-1000-1 |

**ARRIVE Essential 10 Checklist**

| **Item** | **Application** |
|---|---|
| Ethics | Approved by the University of Kentucky IACUC (2018-2968). |
| Sex | Described in each figure. |
| Inclusion criteria | Based on sex, age, body weight, and overt health appearance in each experiment. |
| Exclusion criteria | Based on sex, age, body weight, or medical cases reported by a veterinarian. |
| Sample size | Described in each figure legend. |
| Sample size calculation | None |
| Primary endpoint | Atherosclerosis or renal pathologies |
| Randomization | Study mice were numbered and grouped randomly |
| Blinding | None for experimental design. Some data were measured by two investigators independently to validate the consistency of measurements. |
| Statistical analysis | SigmaPlot 15.0 (SYSTAT Software Inc., CA) or R Statistical Software. |
| Statistical method | Described in each figure legend |
| Data availability | All numerical data used for figures are available in the Supplemental Excel File. |

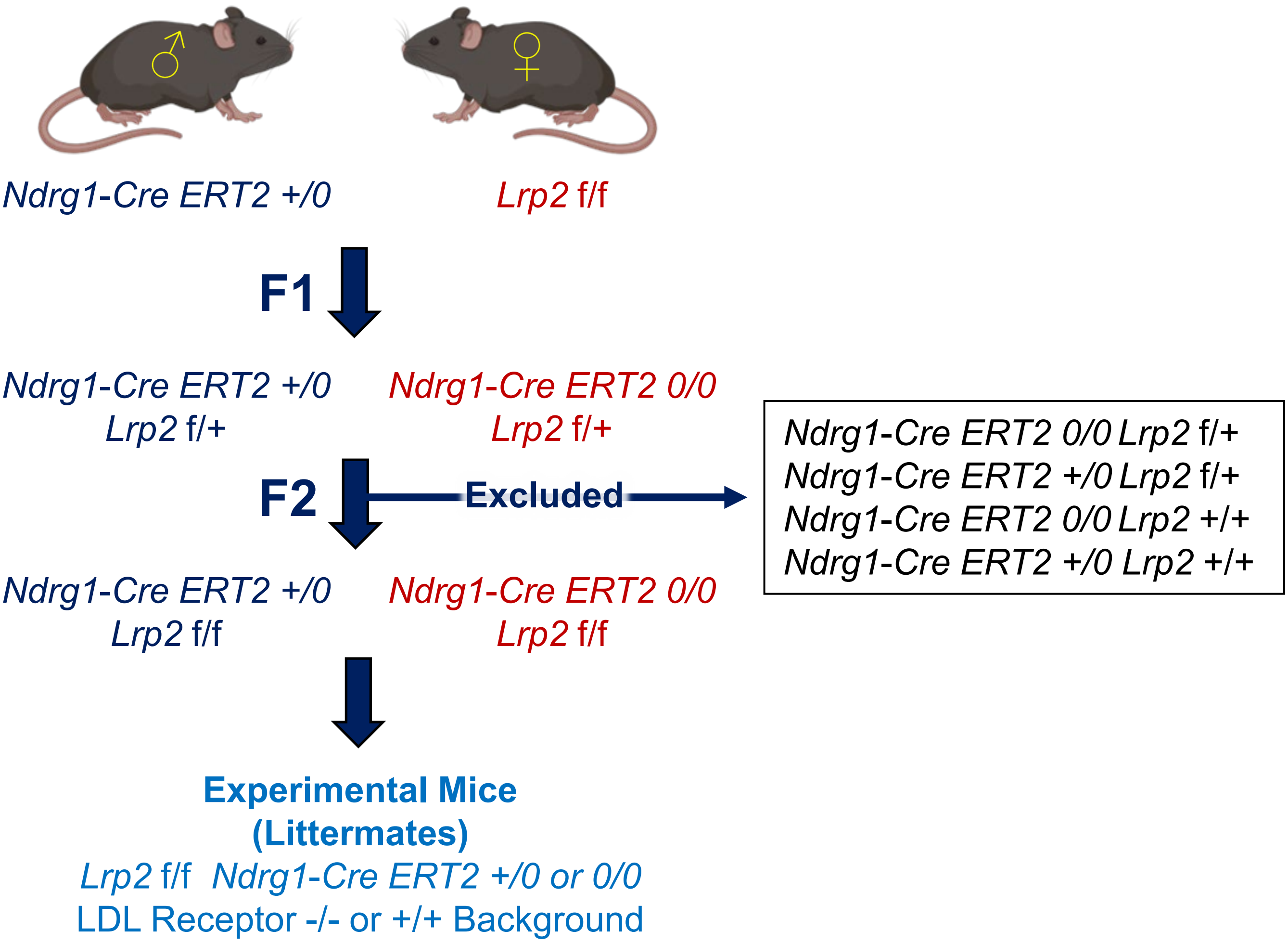


**Figure S1. Breeding strategy to develop inducible PTC-LRP2 +/+ and PTC-LRP2 -/- littermates for experiments.** Study mice were either LDL receptor +/+ or -/- on a C57BL/6J background as indicated in each figure. PTC-specific deletion of megalin was induced by intraperitoneal injection of tamoxifen for 5 consecutive days when they were 4-6 weeks old.

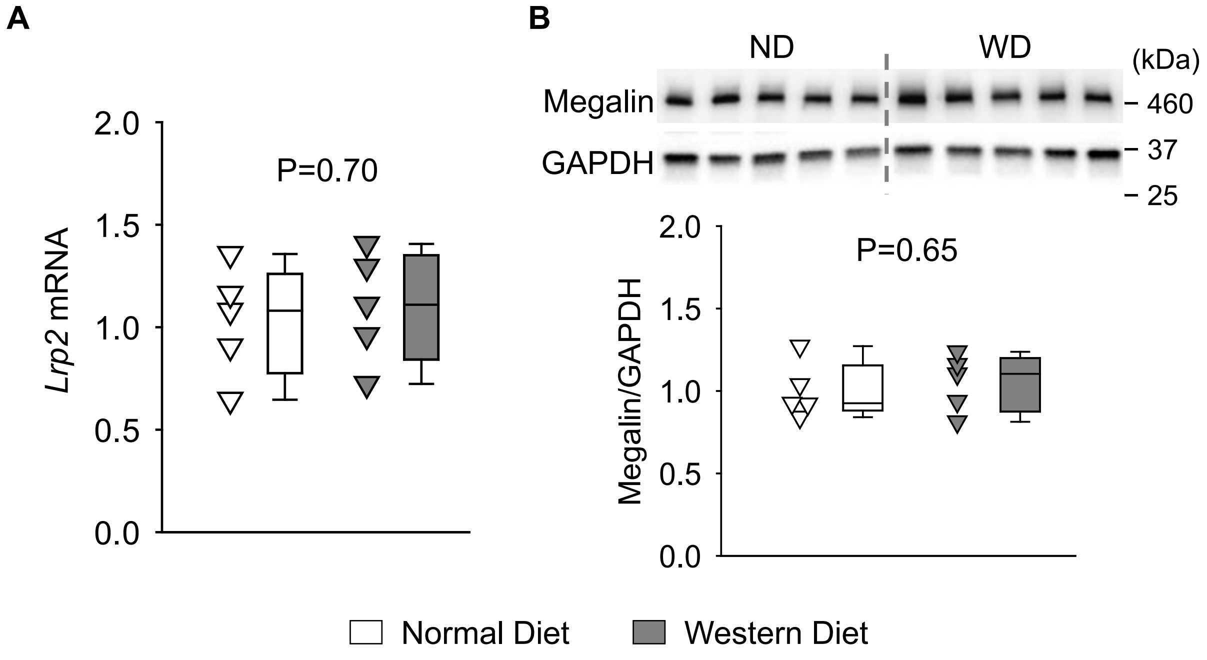


**Figure S2. Western diet feeding did not change renal megalin abundance in mice. (A)** qPCR and **(B)** Western blots of megalin in the kidney of wild-type male mice fed a normal diet versus a Western diet for 12 weeks. N=5/group. P values were calculated by Student's t-test. ND indicates normal diet; WD indicates Western diet.

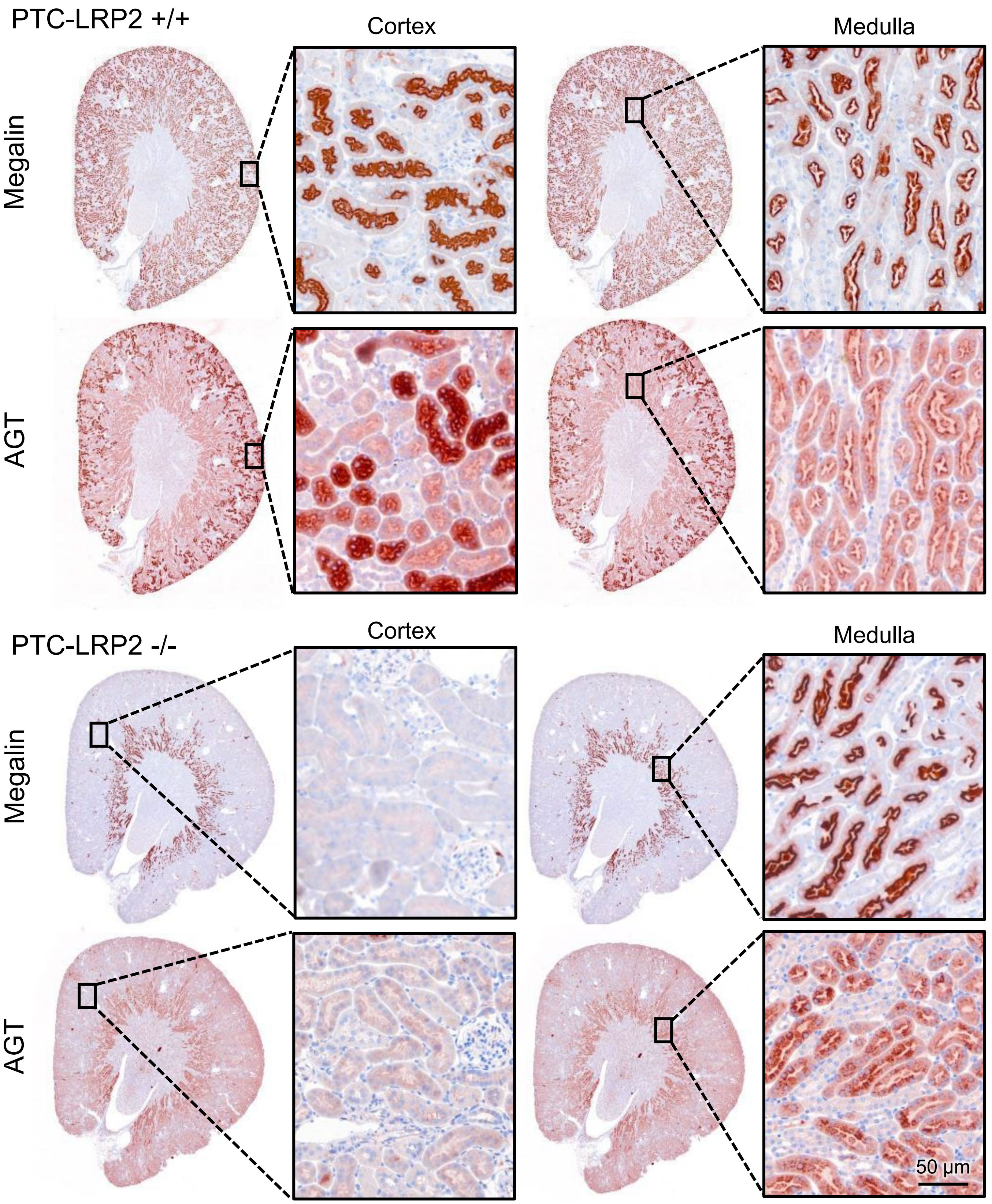


**Figure S3. Immunostaining of megalin and AGT in PTC-LRP2 +/+ and PTC-LRP2 -/- mice.** Immunostaining of megalin or AGT. Four to six-week-old male mice in an LDL receptor -/- background received intraperitoneal injections of tamoxifen for 5 consecutive days. Kidney tissues were harvested and sectioned at 14 weeks after completing the tamoxifen injection.

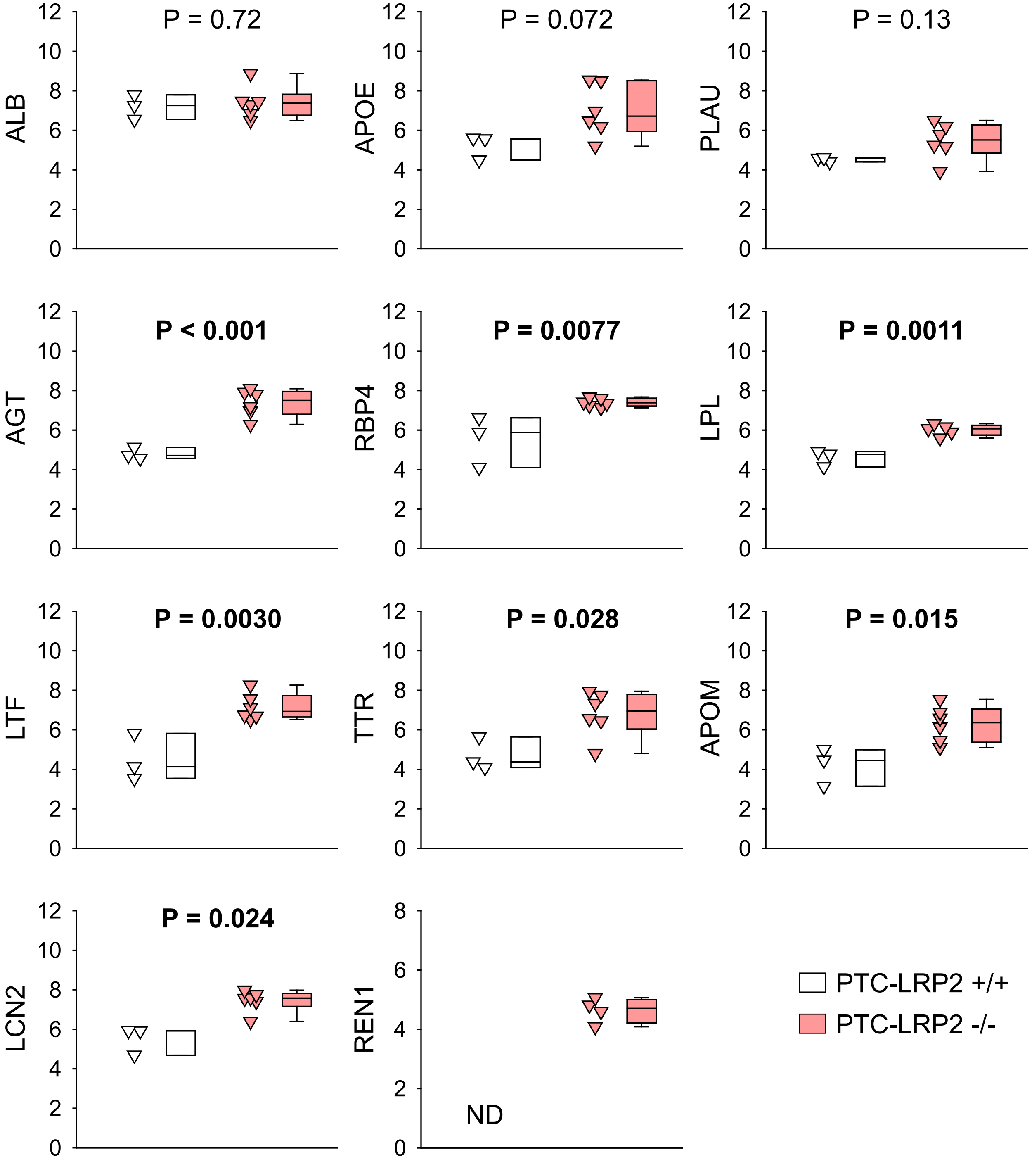


**Figure S4. PTC-specific megalin deletion increased urinary concentrations of multiple megalin ligands.** Urine protein abundance measured by mass spectrometry-assisted proteomics analysis of ALB, APOE, PLAU, AGT, RBP4, LPL, LTF, TTR, APOM, LCN2, REN1 in male PTC-LRP2 +/+ versus -/- mice at 2 weeks after completion of tamoxifen injection. N=3-5/group. P values were calculated by Mann-Whitney test for LCN2 and Student's t-test for other molecules. ND indicates not detected.

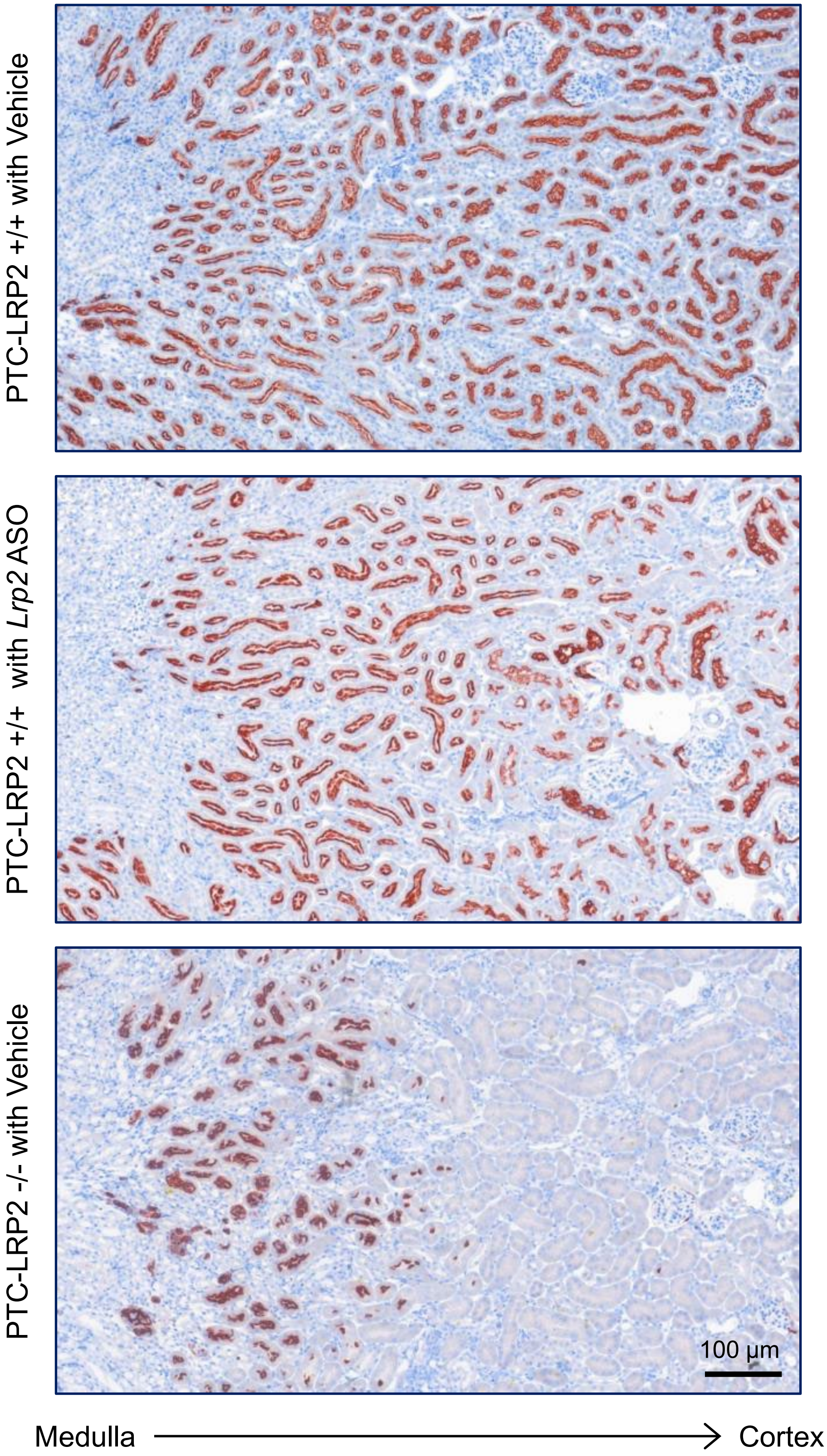


**Figure S5. *Lrp2* ASO did not ablate megalin in proximal convoluted tubules.** Immunostaining of megalin was performed in male PTC-LRP2 +/+ injected with PBS (vehicle) or *Lrp2* ASO and PTC-LRP2 -/- mice injected with vehicle.

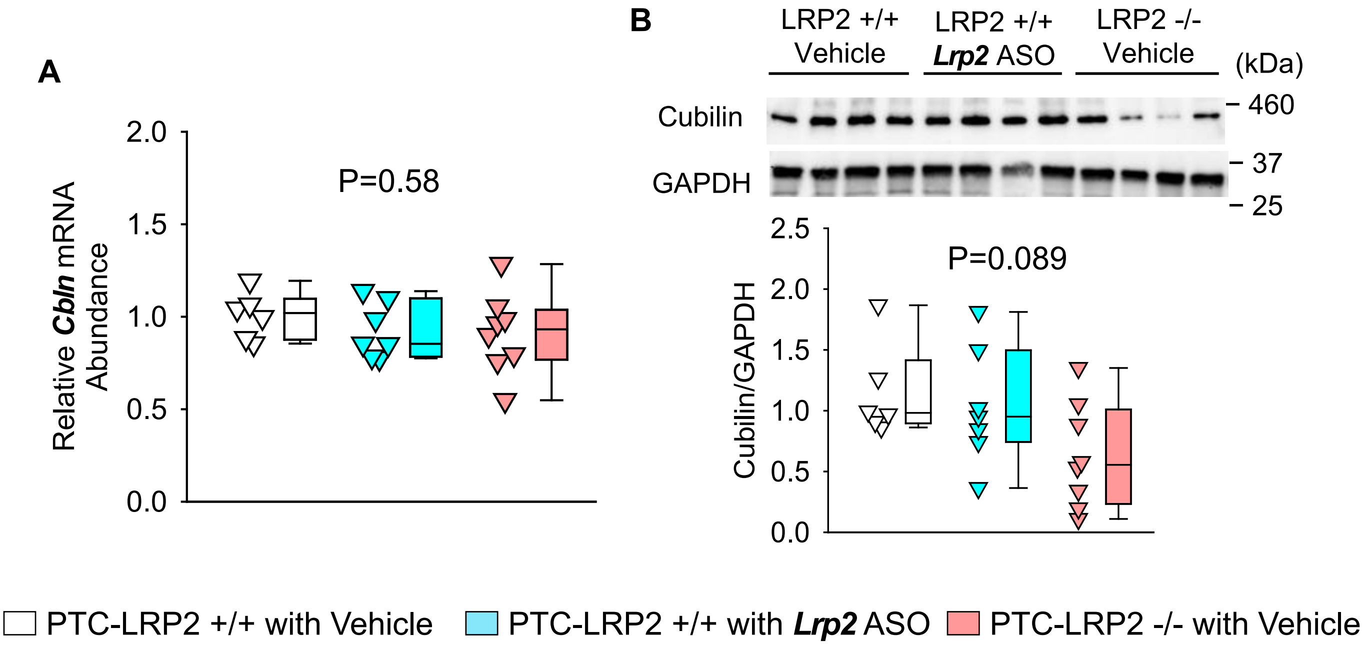


**Figure S6. PTC-specific megalin deletion did not change cubilin abundance. (A)** qPCR and **(B)** Western blot analysis for cubilin in the kidney of male PTC-LRP2 +/+ mice with vehicle injection, PTC-LRP2 +/+ mice with ***Lrp2*** ASO injection, and PTC-LRP2 -/- mice with vehicle injection. Samples were collected from the surface of the kidney. N=8-9/group. P value was calculated by one-way ANOVA.

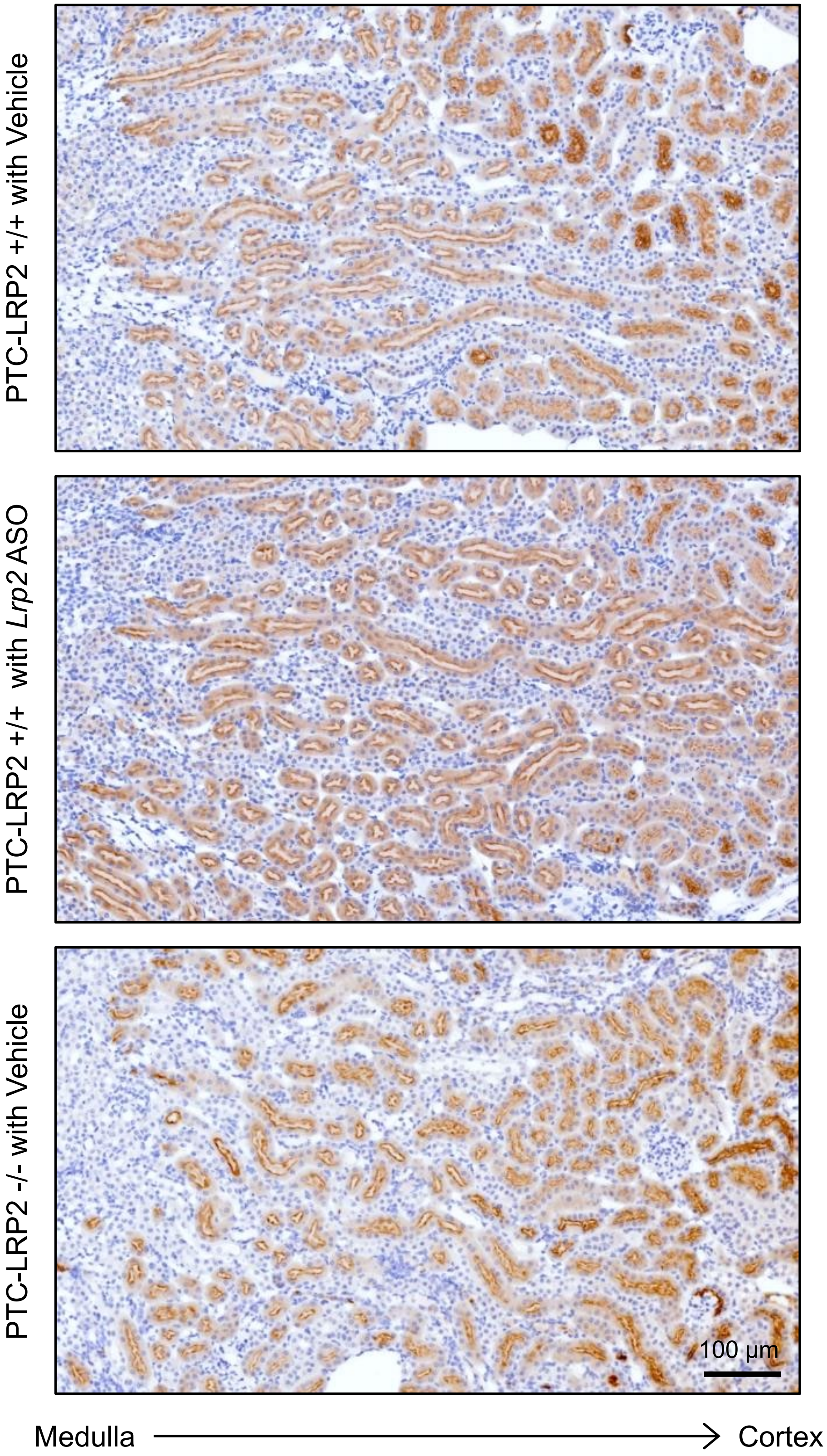


**Figure S7. Inhibition or deletion of megalin did not ablate cubilin in proximal convoluted tubules.** Immunostaining of cubilin was performed in male PTC-LRP2 +/+ injected with PBS (vehicle) or *Lrp2* ASO and PTC-LRP2 -/- mice injected with vehicle.

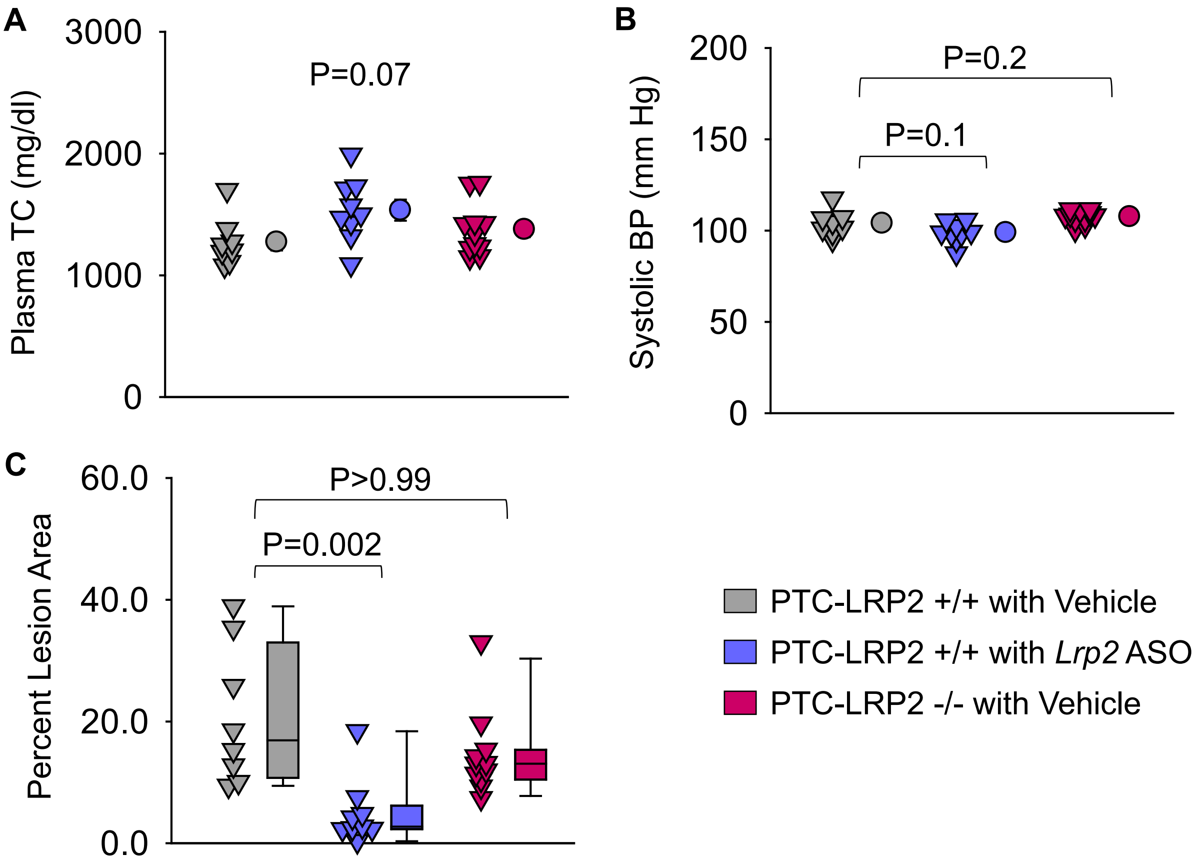


**Figure S8. PTC-specific megalin deletion did not attenuate atherosclerosis in female mice.** Female PTC-LRP2 +/+ and PTC-LRP2 -/- mice on an LDL receptor -/- background were fed a Western diet for 12 weeks. **(A)** Plasma TC (total cholesterol) concentrations were measured using an enzymatic method. **(B)** Systolic BP (blood pressure) was measured using a tail-cuff system. **(C)** Atherosclerosis was measured and quantified using an *en face* method. Statistical analyses: one-way ANOVA followed by the Holm-Sidak test (**A** and **B**) or Kruskal-Wallis one-way ANOVA on ranks followed by the Dunn method (**C**).

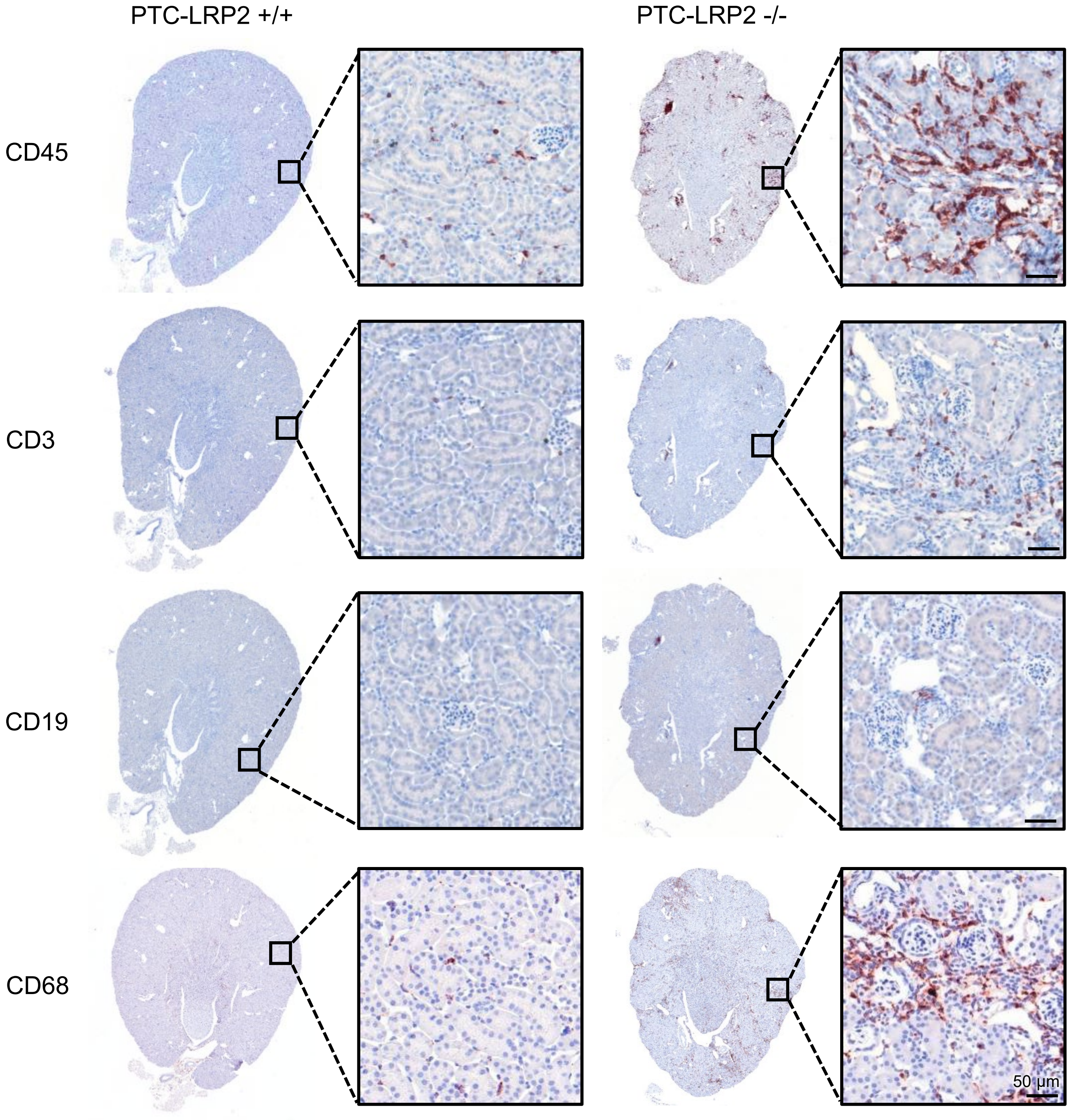


**Figure S9. PTC-specific megalin deletion led to inflammation in male mice fed Western diet.** Four to 6-week-old male mice in an LDL receptor -/- background received intraperitoneal injections of tamoxifen for 5 consecutive days. Two weeks after completing the tamoxifen injection, all study mice were fed a Western diet. Immunostaining of CD45, CD3, CD19, and CD68 was performed on kidney sections of mice fed Western diet for 12 weeks.

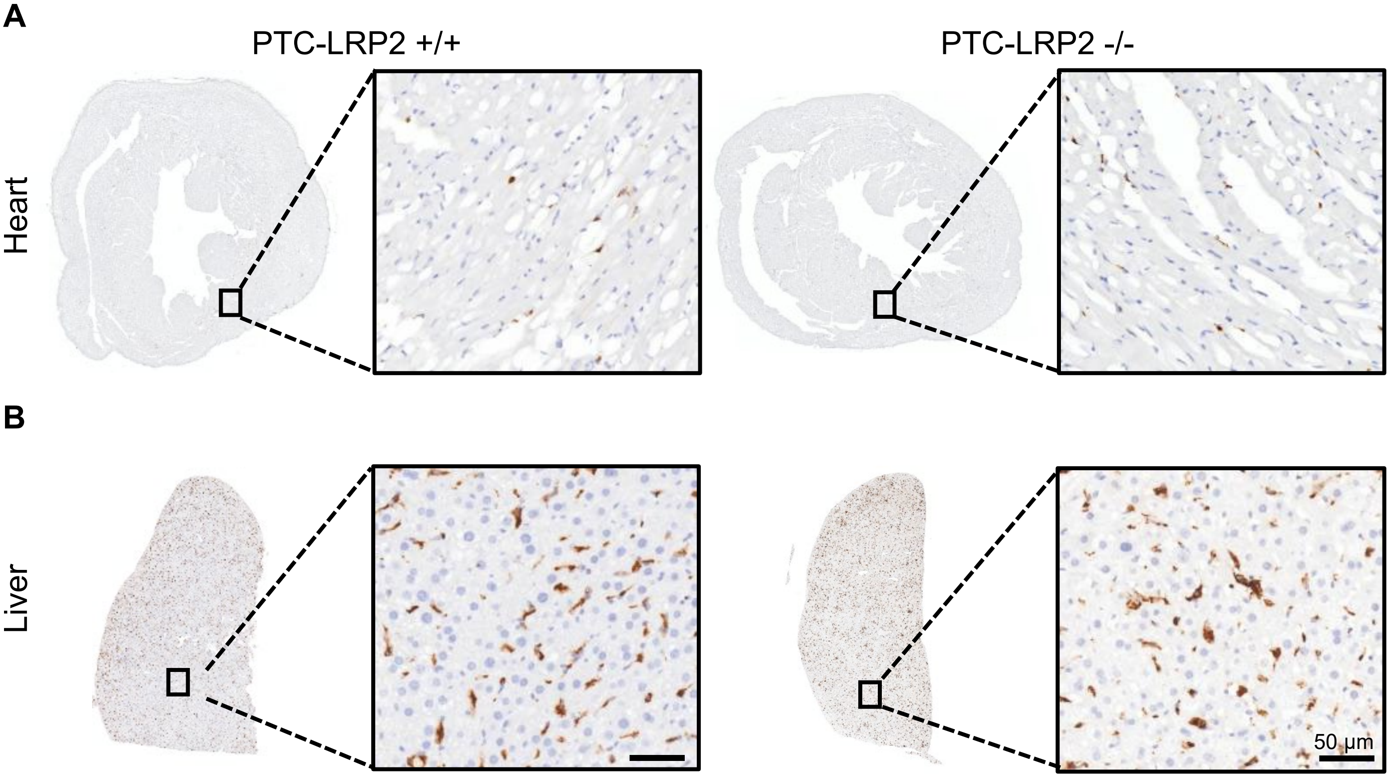


**Figure S10. PTC-specific megalin deletion did not affect macrophage accumulation in the heart and liver of mice fed Western diet for 12 weeks.** Male PTC-LRP2 +/+ versus -/- mice on an LDL-/-background were fed a Western diet. Immunostaining of CD68 in the heart **(A)** and liver **(B**) of PTC-LRP2 +/+ versus -/- mice fed a Western diet for 12 weeks.

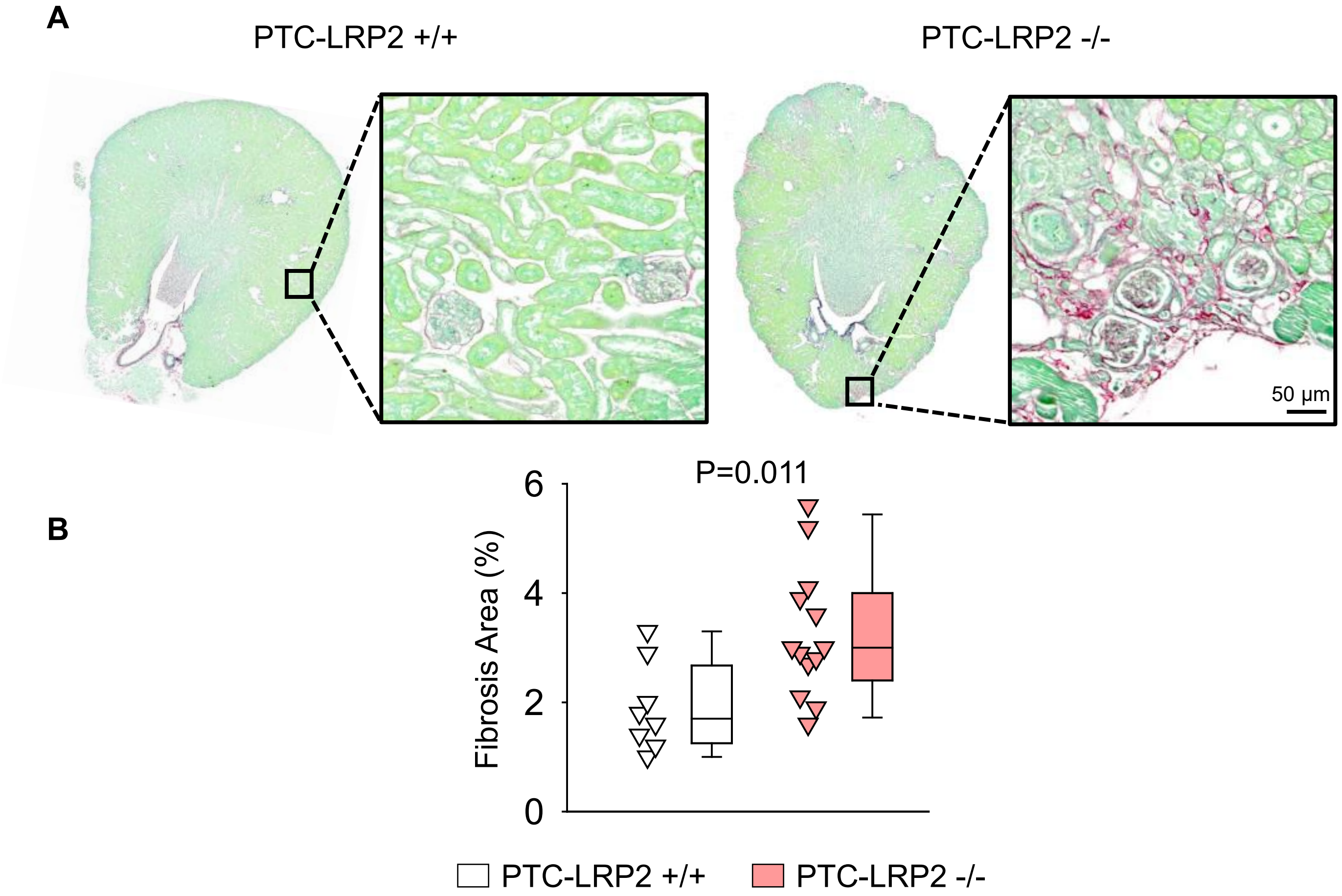


**Figure S11. PTC-specific megalin deletion led to renal fibrosis in male mice fed Western diet.** Representative images of picrosirius red staining and its quantification in the kidney of male PTC-LRP2 +/+ versus -/- mice fed Western diet for 12 weeks. N=8-13/group. Statistical analysis: Student's t-test.

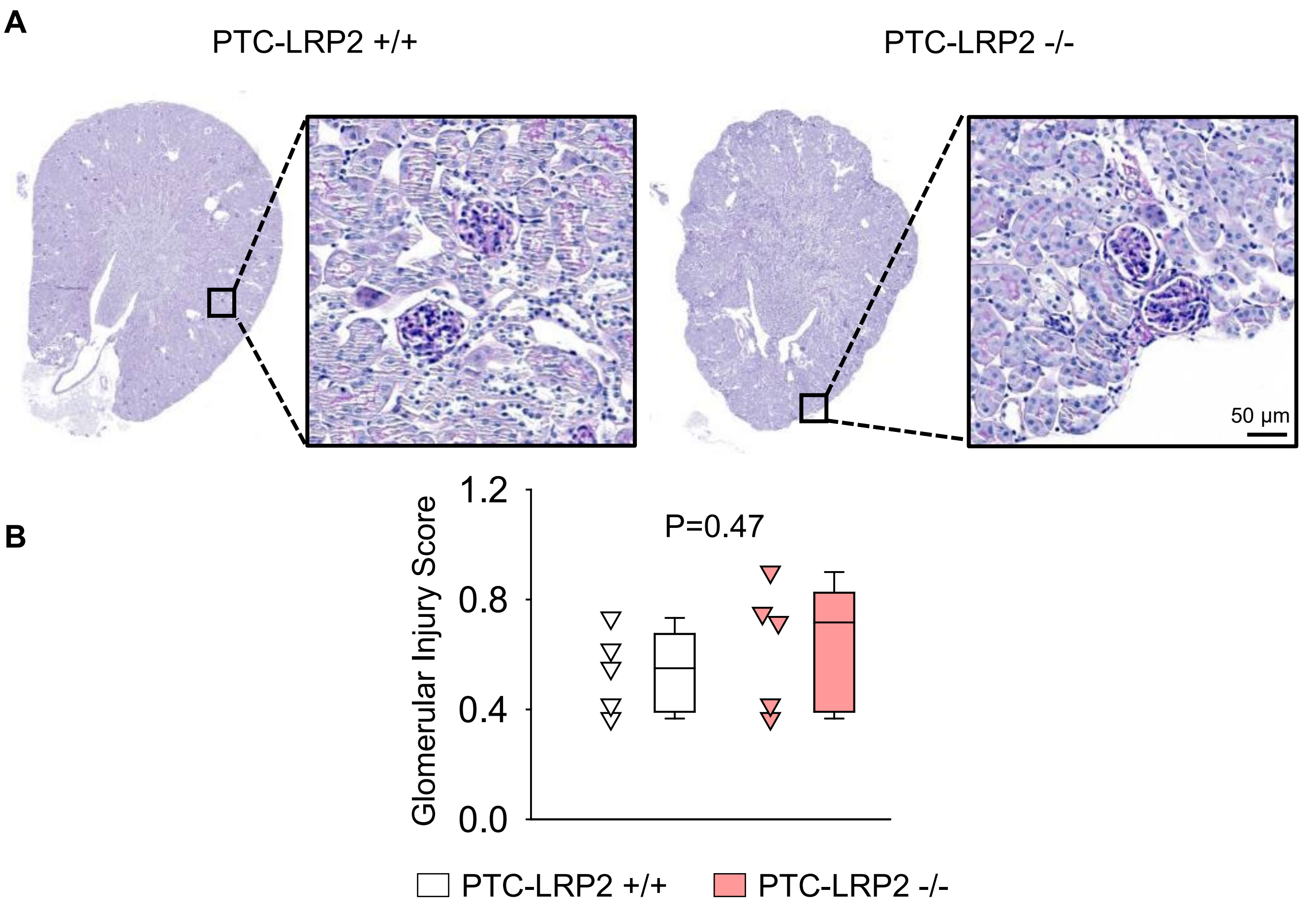


**Figure S12. PTC-specific megalin deletion did not lead to glomerular damage in male mice fed Western diet.** Representative images of PAS staining and the quantification of glomerular damage in the kidney of male PTC-LRP2 +/+ versus -/- mice fed Western diet for 12 weeks. N=5/group. Statistical analysis: Student's t-test.

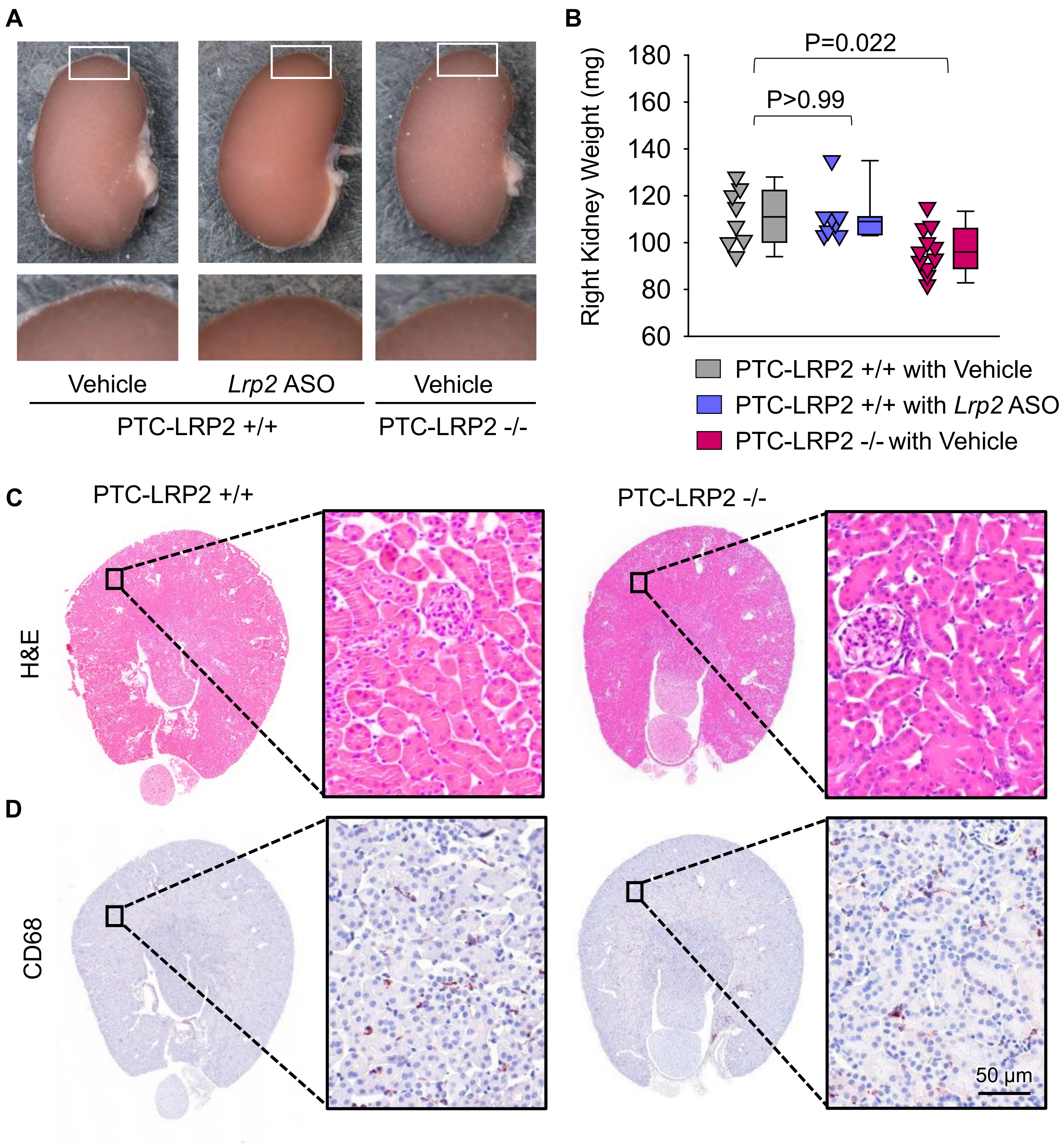


**Figure S13. PTC-specific megalin deletion did not cause significant pathology in female LDL receptor -/- mice.** Female PTC-LRP2 +/+ versus -/- mice in an LDL receptor -/- background were fed a Western diet for 12 weeks. Gross images of kidneys **(A)** and kidney weight **(B)** at termination. **(D)** H&E staining and **(E)** immunostaining of megalin in cross-sections of mouse kidney. Statistical analysis: Kruskal–Wallis one-way ANOVA on Ranks test followed by Dunn's test (**B**).

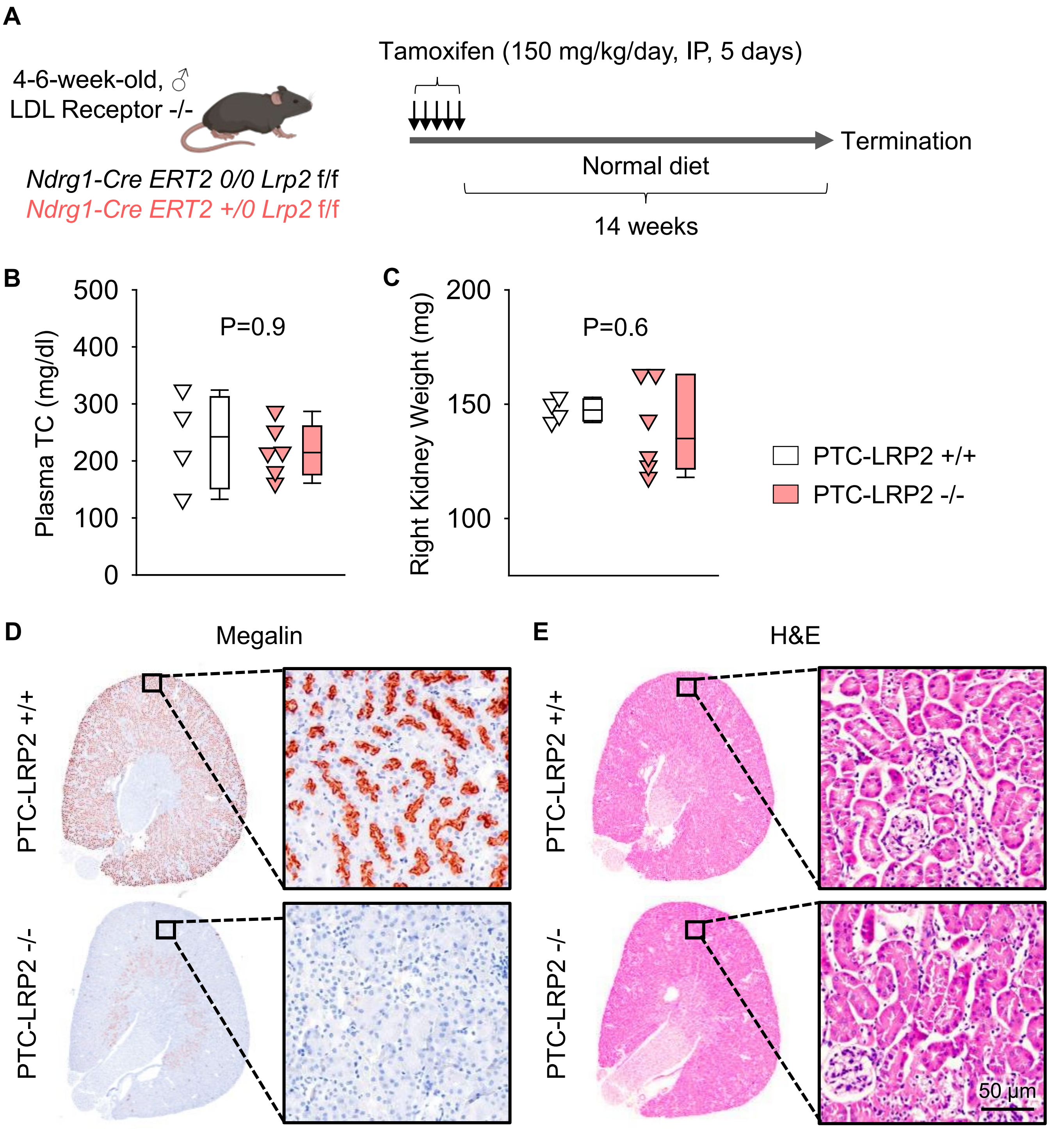


**Figure S14. PTC-specific megalin deletion did not cause significant renal pathology in male LDL receptor -/- mice fed a normal diet. (A)** Male PTC-LRP2 +/+ versus -/- mice in an LDL receptor -/- background were fed a normal laboratory rodent diet for 15 weeks after tamoxifen induction. **(B)** Plasma TC (total cholesterol) concentrations were measured using an enzymatic method. **(C)** Kidney weight was measured at termination. **(D)** Immunostaining of megalin in cross-sections of mouse kidneys. **(E)** H&E staining in cross-sections of mouse kidney. Statistical analysis: Mann-Whitney U-test (**B** and **C**).

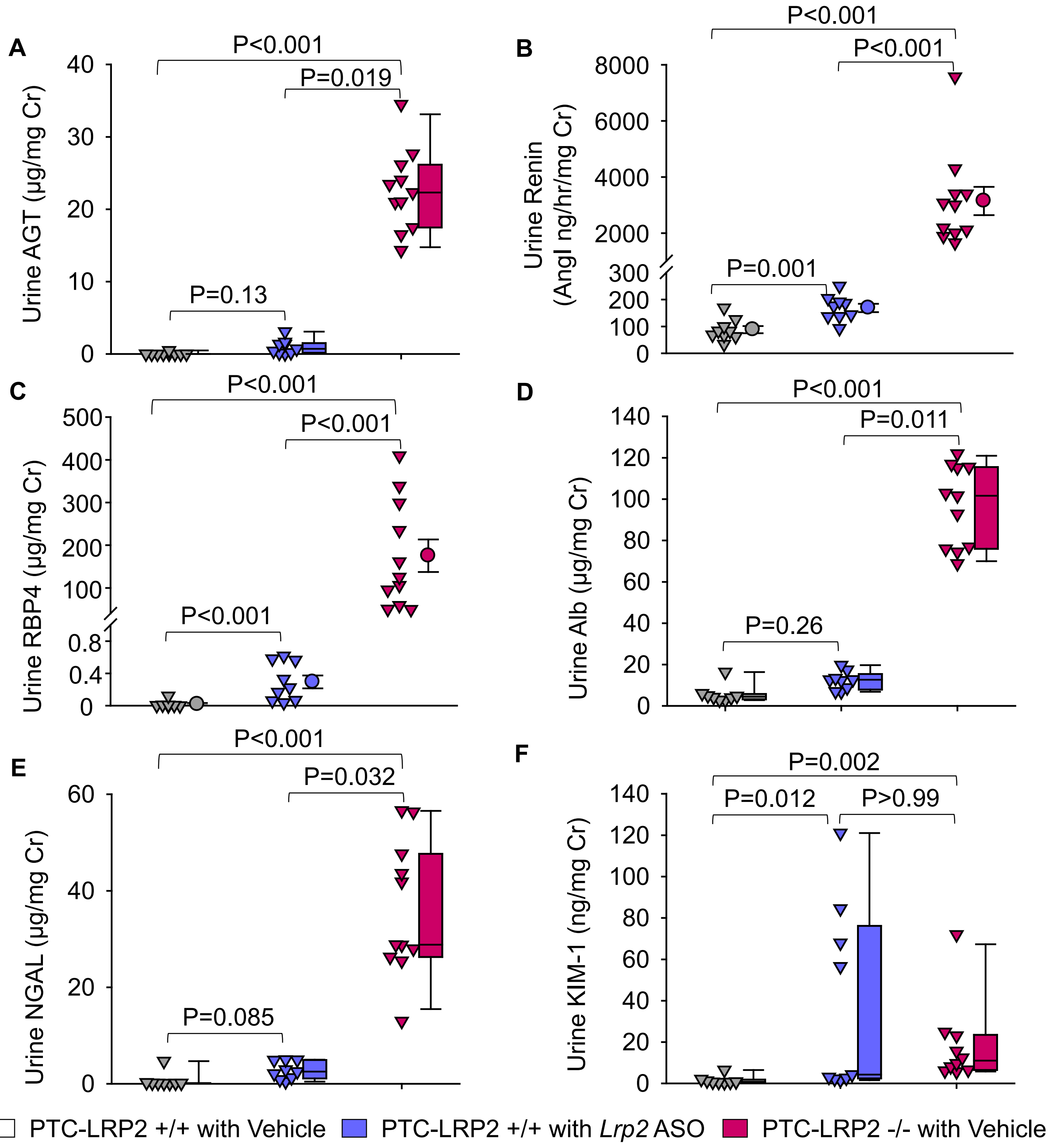


**Figure S15. PTC-specific megalin deletion increased renal PTC injury markers in female mice.** Four to 6-week-old female mice in an LDL receptor -/- background received intraperitoneal injections of tamoxifen for 5 consecutive days. Two weeks after completing the tamoxifen injection, all study mice were fed a Western diet for 12 weeks. The study mice received PBS (Vehicle) or *Lrp2* antisense oligonucleotides (*Lrp2* ASO, 6 mg/kg/week) injection started 1 week prior to Western diet feeding. Urine was collected before termination. AGT **(A)**, renin **(B)**, RBP4 **(C)**, albumin **(D)**, NGAL **(E)**, and KIM-1 **(F)** in urine were measured using ELISA kits and normalized by urine creatinine concentrations. Statistical analysis: Kruskal-Wallis one-way ANOVA on Ranks with Dunn post hoc test **(A, D-F)** or one-way ANOVA with Holm-Sidak post hoc test **(B, C)**.

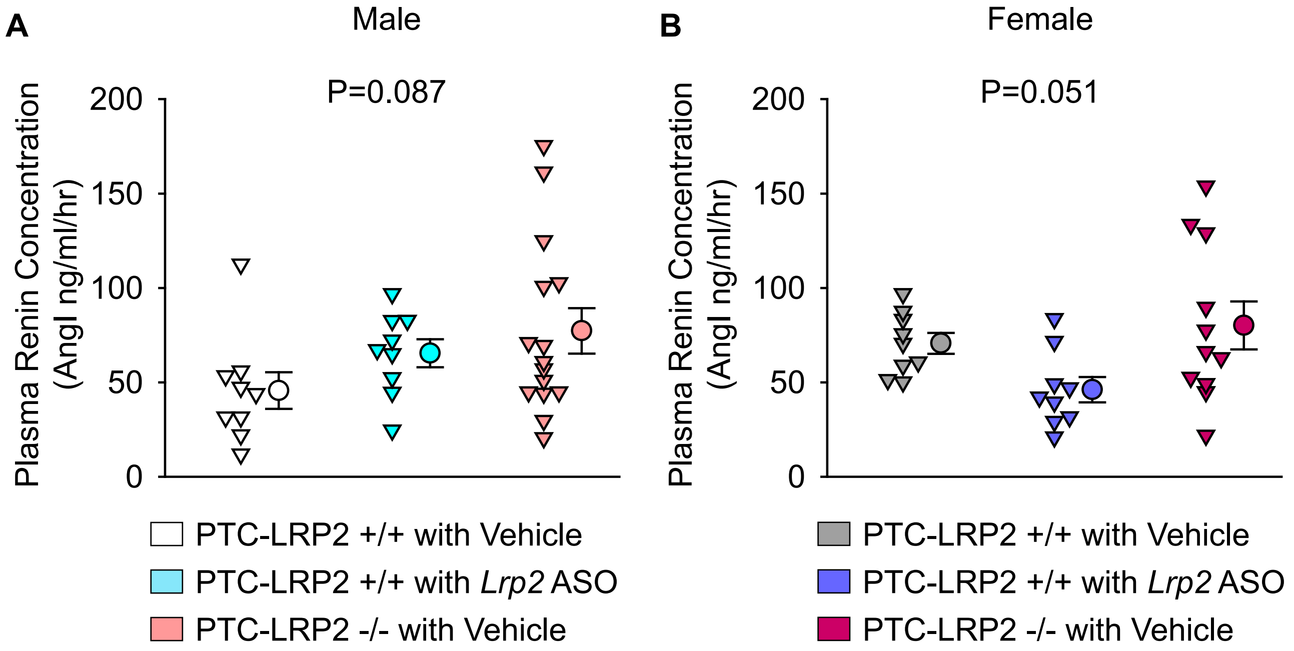


**Figure S16. PTC-specific megalin deletion did not change plasma renin concentrations in both male and female mice.** Male or female PTC-LRP2 +/+ versus -/- mice on an LDL receptor -/- background were fed a Western diet for 12 weeks. Plasma renin concentrations were measured using an ELISA method in male **(A)** and female **(B)** mice. Statistical analyses: One-way ANOVA.

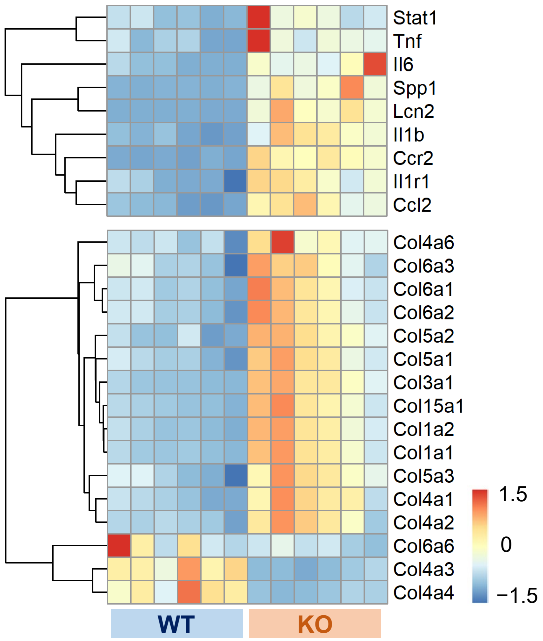


**Figure S17. PTC-specific megalin deficiency augmented inflammation-related transcriptomes in male mice after 12 weeks of Western diet.** Heatmap with Z-scored coloring displaying DEGs associated with inflammation and collagens at 12 weeks of Western diet. N=6/group.

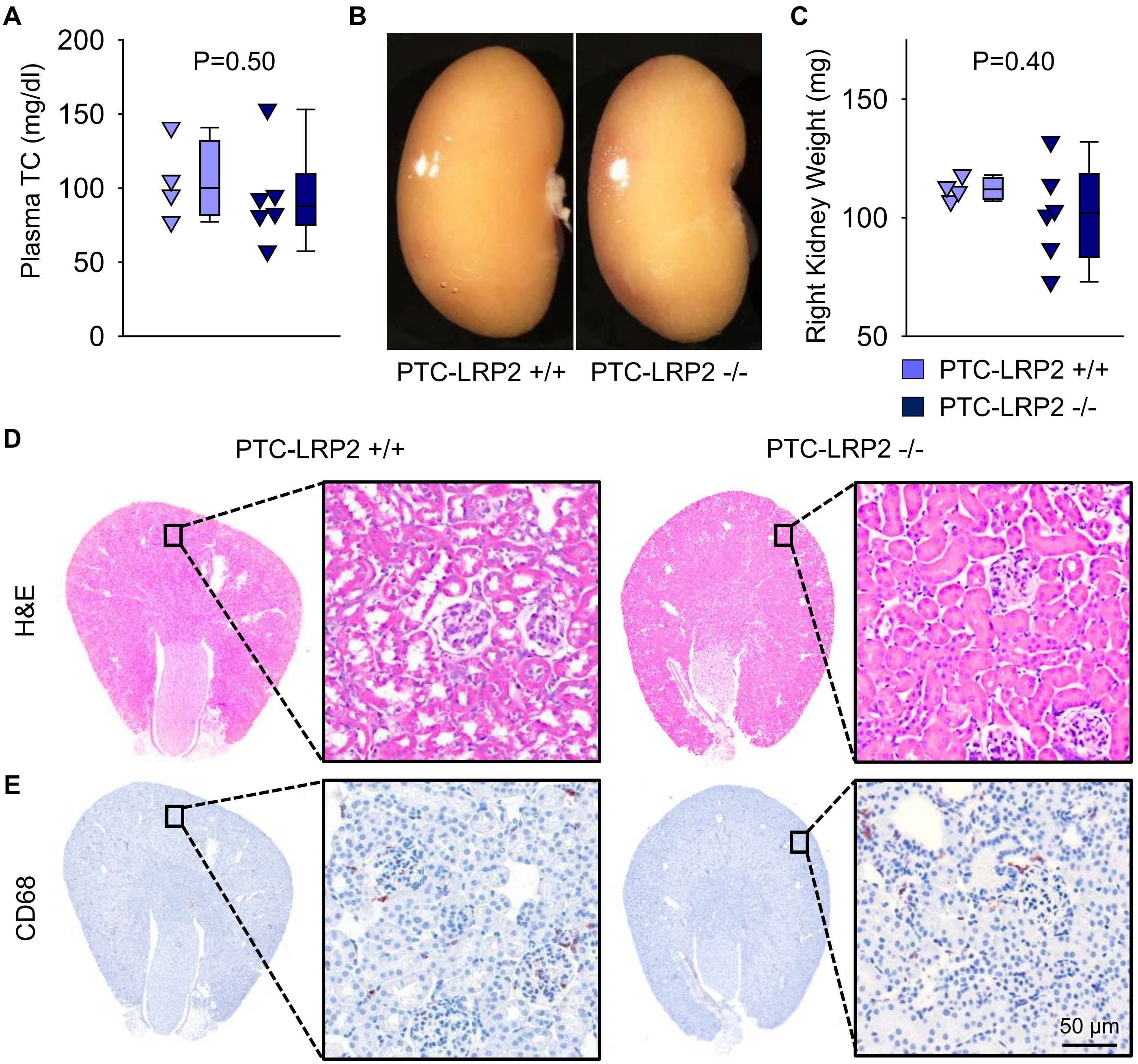


**Figure S18. PTC-specific megalin deletion did not cause significant pathological changes in female LDL receptor +/+ mice.** Female PTC-LRP2 +/+ and PTC-LRP2 -/- mice on a C57BL/6J background were fed a Western diet for 12 weeks. **(A)** Plasma TC (total cholesterol) concentrations were measured using an enzymatic method. Gross images of kidneys **(B)** and kidney weight **(C)** at termination. **(D)** H&E staining and **(E)** immunostaining of CD68 in tissue sections of mouse kidneys. Statistical analysis: Mann-Whitney U-test (**A** and **C**).

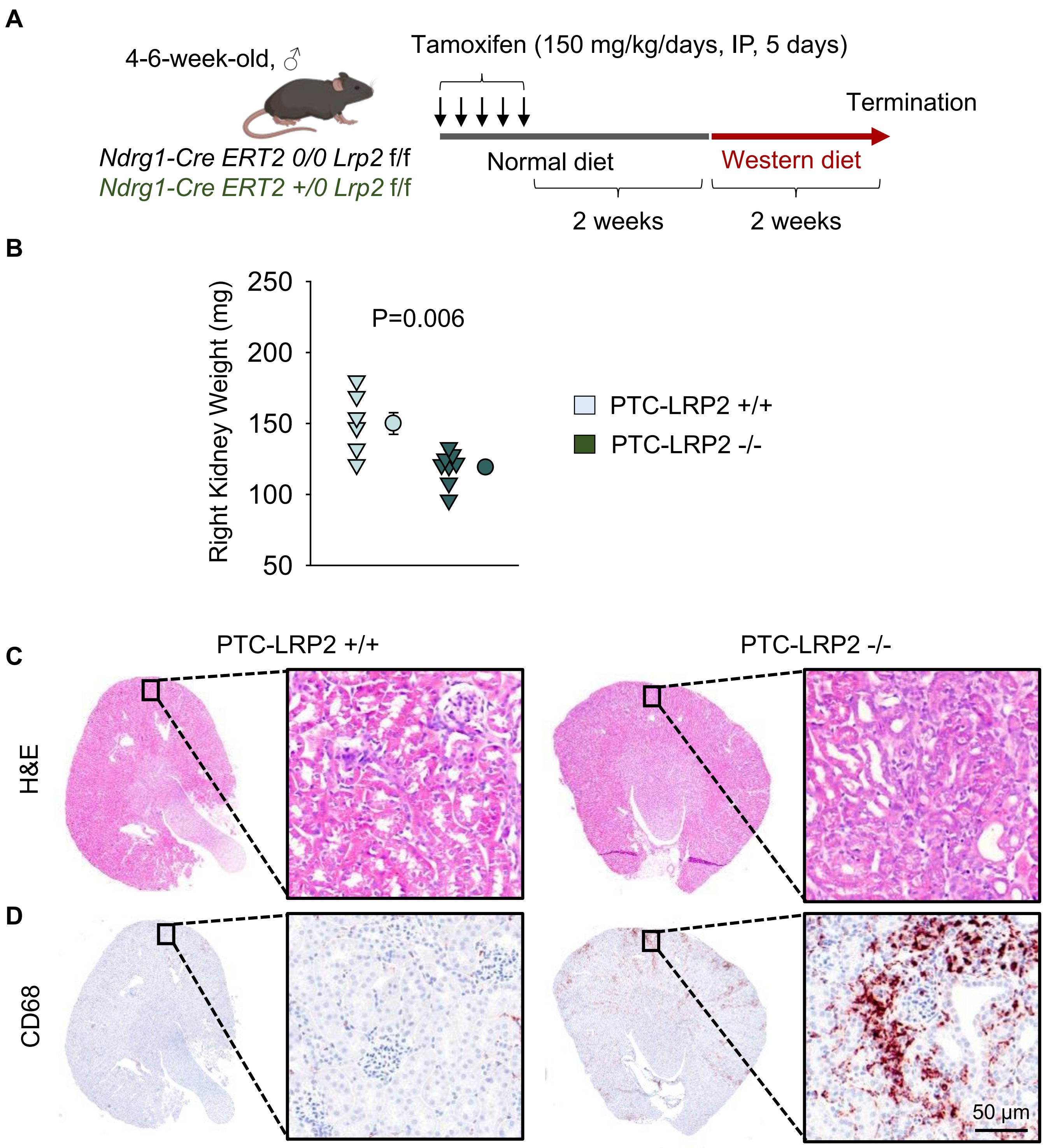


**Figure S19. PTC-specific megalin deletion caused renal pathologies in male LDL receptor +/+ mice.** Male PTC-LRP2 +/+ and PTC-LRP2 -/- mice on a C57BL/6J background were fed a Western diet for 2 weeks **(A)**. **(B)** Kidney weight was measured at termination and analyzed by Student's t-test. **(C)** H& E staining and **(D)** immunostaining of CD68 in cross-sections of mouse kidneys.
